# Mitochondrial regulator PGC-1a in neuronal metabolism and brain aging

**DOI:** 10.1101/2023.09.29.559526

**Authors:** Dylan C Souder, Eric R McGregor, Timothy W Rhoads, Josef P Clark, Tiaira J Porter, Kevin Eliceiri, Darcie L Moore, Luigi Puglielli, Rozalyn M Anderson

**Author notes:** equal contributors (Dylan C Souder, Eric R McGregor).

## Abstract

The brain is a high energy tissue, and the cell types of which it is comprised are distinct in function and in metabolic requirements. The transcriptional co-activator PGC-1a is a master regulator of mitochondrial function and is highly expressed in the brain; however, its cell-type specific role in regulating metabolism has not been well established. Here, we show that PGC-1a is responsive to aging and that expression of the neuron specific PGC-1a isoform allows for specialization in metabolic adaptation. Transcriptional profiles of the cortex from male mice show an impact of age on immune, inflammatory, and neuronal functional pathways and a highly integrated metabolic response that is associated with decreased expression of PGC-1a. Proteomic analysis confirms age-related changes in metabolism and further shows changes in ribosomal and RNA splicing pathways. We show that neurons express a specialized PGC-1a isoform that becomes active during differentiation from stem cells and is further induced during the maturation of isolated neurons. Neuronal but not astrocyte PGC-1a responds robustly to inhibition of the growth sensitive kinase GSK3b, where the brain specific promoter driven dominant isoform is repressed. The GSK3b inhibitor lithium broadly reprograms metabolism and growth signaling, including significantly lower expression of mitochondrial and ribosomal pathway genes and suppression of growth signaling, which are linked to changes in mitochondrial function and neuronal outgrowth. In vivo, lithium treatment significantly changes the expression of genes involved in cortical growth, endocrine, and circadian pathways. These data place the GSK3b/PGC-1a axis centrally in a growth and metabolism network that is directly relevant to brain aging.

## INTRODUCTION

“Mitochondrial dysfunction” has been proposed as a hallmark of the aging process (López-Otín et al., 2023), and changes in mitochondrial pathways are a key feature of calorie restriction (CR) (Barger et al., 2015), a dietary intervention that can extend lifespan in diverse species. In all cell types, the response to regulatory inputs or changes in the environment requires the ability to divert metabolites to the appropriate synthetic pathways while meeting energetic demand. In this way, the integrity of mitochondrial sensing and adaptation is fundamental to cellular resilience and functionality (Monzel et al., 2023). Multiple lines of evidence now point to a role for mitochondria in age-related disease and mitochondrial dysfunction has been linked to disorders as different as cancer, cardiovascular disease, and neurodegeneration (Harrington et al., 2023; Lima et al., 2022). The ubiquitously expressed transcriptional co-activator peroxisome proliferator-activated receptor gamma-coactivator 1 alpha (PGC-1a) has been described as the master regulator of mitochondrial function. PGC-1a plays a key role in integrating and responding to diverse signals, ensuring the cellular metabolic response matches the prevailing conditions (Miller, Clark, & Anderson, 2019). In cultured cells, modest PGC-1a overexpression leads not only to mitochondrial activation but extends to extra-mitochondrial processes such as redox and lipid homeostasis, chromatin maintenance, cell cycle, growth, and structural remodeling (Miller, Clark, Martin, et al., 2019). Despite its key role in mitochondrial adaptation, the regulation of PGC-1a outside of the liver, skeletal muscle, and adipose tissue remains poorly defined, and this is particularly true for the brain. Failure in maintaining brain energetics has been clearly linked to neurodegenerative disorders (Cunnane et al., 2020), and yet cell-type specific details of energy balance regulation and a precise role for PGC-1a have not been defined.

The activity of the PGC-1a protein is controlled by numerous upstream regulators, post-translational modifications, and cell-type and context-specific binding partners that fine-tune its co-activator function (Miller et al., 2019a). There are multiple isoforms of PGC-1a derived from distinct promoter regions of the *Ppargc-1a* gene (Martínez-Redondo et al., 2015). The promoters are activated in response to different physiological stimuli (Ruas et al., 2012; Soyal et al., 2020), and the resulting protein isoforms appear to have both common and isoform specific gene targets (Lozoya et al., 2021; Martínez-Redondo et al., 2015; Ruas et al., 2012). For example, in skeletal muscle, PGC-1a transcript variants expressed off the alternative promoter are selectively induced by resistance exercise, while the canonical promoter is activated by endurance exercise (Ruas et al., 2012). These differences in promoter activity also extend to tissue-specific differences in PGC-1a isoform expression, as skeletal muscle, white adipose tissue, and liver display unique profiles of PGC-1a isoform abundance (Martinez-Redondo et al., 2015). In the central nervous system, a third promoter >500kb upstream of the canonical transcription start site is active and has been shown to be responsive to hypoxia (Soyal et al., 2020; Soyal, Felder, Auer, Hahne, Oberkofler, Witting, Paulmichl, Landwehrmeyer, Weydt, & Patsch, 2012). It is as yet unclear how functionally redundant the PGC-1a isoforms derived from the brain-specific promoter are with those derived from the canonical and alternate promoters, whether there is specificity in how they are regulated, and the relative abundance among cell types might differ in the brain.

Although not validated in primary cells, we previously reported that glycogen synthase-kinase 3-beta (GSK3b) regulates mitochondrial energy metabolism via PGC-1a in glioma cells (H4) and in phaeochromocytoma-differentiated neurons (PC12) (Martin et al., 2018). GSK3b inhibition increased mitochondrial respiration and membrane potential and altered NAD(P)H metabolism, and these changes were associated with activation of Ppargc1a canonical and alternate promoters. Our previous studies in mice indicated that aging impacts mitochondrial and redox metabolism in the hippocampus in a region and cell-type specific manner and that these changes are linked to lower PGC-1a protein levels (Martin et al., 2016). GSK3b and PGC-1a show the same pattern of expression in the hippocampus in mice and in monkeys, and their relative abundance is reflected in the level of mitochondrial activity. Here we focus on aging to determine the impact of aging on metabolism and on PGC-1a. We investigate the role of PGC-1a in primary neurons and astrocytes and determine the extent to which GSK3b regulates PGC-1a expression driven from the three different promoters. We define the impact of GSK3b inhibition on neuronal metabolism and growth indices and validate the conservation of these effects in vivo. Together these experiments make the case that the PGC-1a plays a role in brain aging and that differential PGC-1a isoform expression and differences in GSK3b sensitivity allow for distinct regulation of metabolism among brain resident cell types.

## RESULTS

### Aging impacts brain immune, inflammatory, and neuronal functional networks

To investigate aging across the lifespan, C3B6-F1 hybrid male mice were maintained on a fixed calorie intake to avoid obesity and to maximize health during aging (Martin et al., 2016; Miller et al., 2017). Three age groups were defined: adult (10 months), late-middle age (20 months), and advanced age (30 months). To gain a molecular perspective on brain aging, RNA sequencing was conducted on brain hemispheres (n = 5 per age group), resulting in 1.6 billion sequencing reads (∼105 million per sample). After trimming, reads were aligned to the mouse genome, GRCm39, and approximately 48,000 transcripts from 18,000 genes were detected, identified, and quantified. Differential expression analysis among the three age groups identified age-sensitive genes across the adult lifespan (10 months vs. 30 months; 471 genes), including early (10 months vs. 20 months; 69 genes) and late (20 months vs. 30 months; 47 genes) life-stages (Fig.1A; Fig.S1A; Table S1). Gene Set Enrichment Analysis (GSEA) (Subramanian et al., 2005) analysis was conducted to identify functional pathways impacted by age (Table S1). Aggregate data of age-responsive pathways are shown in the rank plot (Fig.1B, Fig.S1B). Upregulated pathways included immune/ inflammatory (antigen processing, viral infection pathways, JAK/STAT, TLR (Toll-Like Receptor) signaling), proteome maintenance (phagosome, ribosome), and metabolic pathways (type I diabetes, oxidative phosphorylation, lipid, and atherosclerosis). Down-regulated pathways included primarily neuronal functional pathways (neuroactive ligand-receptor interactions, hippo signaling, morphine addiction).

**Figure 1:**
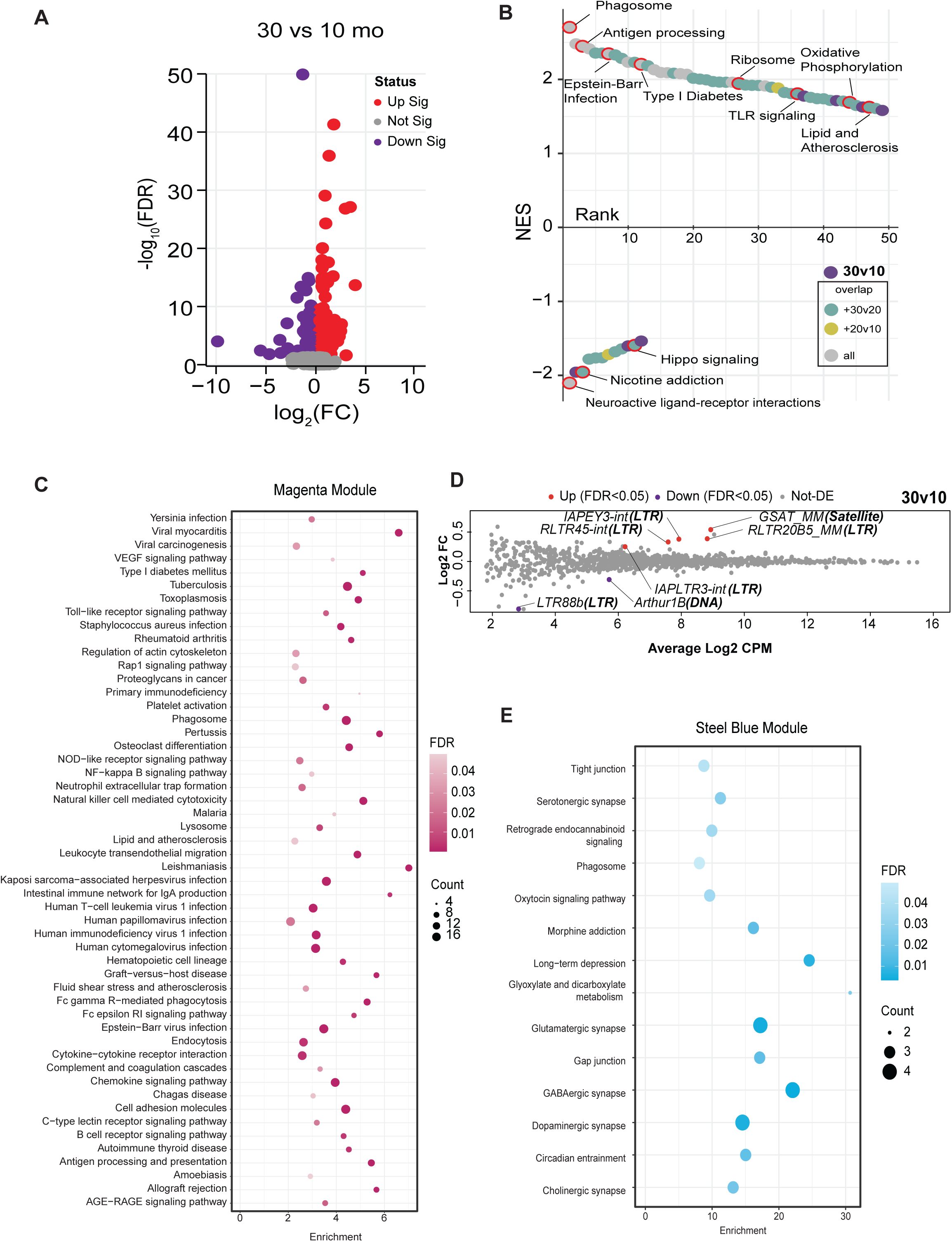
Aging impacts brain immune, inflammatory, and neuronal functional networks. **A)** Volcano plot displaying transcripts quantified. Statistically significant transcripts are highlighted in red (upregulated) or purple (downregulated) in 30-month-old compared to 10-month-old male cortical tissue. **B)** Rank order plot of enriched pathways detected by GSEA. Plot is ranked by normalized enrichment score. Each point represents pathways enriched in the 30v10 comparison. Points are color-coded by overlap with other comparisons (20v10 – yellow, 30v20 – cyan, 30v10 only – purple, present in all – grey). **C)** KEGG pathways enriched in the Magenta module of weighted gene correlation network analysis (WGCNA). **D)** Mean-difference (MD) plots of transposable elements (TE) expression Log2 FC against the average Log2 count-per-million (CPM) for the 30m/10m comparison. TE callouts list the: “TE name (**TE class**).” **E)** KEGG pathways enriched in the Steel blue module of WGCNA.

Weighted gene correlation network analysis (WGCNA) (Langfelder & Horvath, 2008) is an alternate approach that considers the entire transcriptome and identifies clusters of transcripts with a shared pattern of expression. WGCNA identified 30 modules (Table S2) that were then subject to pathway analysis via the Kyoto Encyclopedia of Genes and Genomes (KEGG). Of the 3 modules with high pathway-to-transcript ratios, the largest included immune and inflammatory pathways (magenta module, 467 transcripts, 42 pathways). Notwithstanding redundancy among pathways, the module was highly enriched for viral response, infectious response, B cell, T cell, and complement pathways (Fig.1C). Other pathways of interest included phagosome, lysosome, and endocytosis. The mice were maintained in Specific Pathogen Free conditions indicating that the enrichment of immune and inflammatory pathways was not due to infection but more likely a reflection of immune deregulation and sterile inflammation. The age-related induction of viral infection pathways identified in the GSEA suggested that perhaps native DNA might be triggering the response, in particular since transposable elements (TEs) have been linked to age-associated diseases (Gorbunova et al., 2021). Of the 1146 TE transcripts detected in the RNA-Seq data set (Supplemental Table 3), 7 were differentially expressed at advanced age, (Fig.1D), with 3 of these showing significant differences in expression by late middle-age (Fig.S2). The second module of interest (steel blue module, 41 transcripts, 14 pathways) featured pathways that are integral to neuronal function (Fig.1E). Glutaminergic, dopaminergic, cholinergic, and serotonergic synapse pathways were enriched in this module, along with other pathways involved in intercellular communication such as tight junction and gap junction. The identification of immune, inflammatory, and neurological pathways responsive to aging and age-related conditions is consistent with prior studies (Walker et al., 2022; Wilson et al., 2023) and indicates that the environment of the aging brain is distinct from that of mature adult mice and that processes involved in neuronal function may become compromised with age.

### Metabolism of brain aging is reflected in the transcriptome and proteome

Our prior studies using this same cohort of mice focused specifically on the hippocampus and identified age-related changes in mitochondrial activity and NAD(P)H redox metabolism that were life-stage, region, and cell-type specific (Martin et al., 2016). Applying KEGG analysis to the third module identified via WGCNA (green-yellow module, 432 transcripts, 24 pathways) revealed an enrichment of metabolic pathways (Fig.2A Table S2). Pathways enriched in this module include pyruvate metabolism, glycolysis, Krebs cycle, oxidative phosphorylation, and amino acid metabolism. Neurodegenerative disease pathways are also featured, including Alzheimer’s disease, Parkinson’s disease, and amyotrophic lateral sclerosis, along with autophagy and mitophagy pathways. The green-yellow module was significantly enriched for genes encoding mitochondrial proteins (Fig.2B), with 11% of transcripts overlapping with the MitoCarta (Rath et al., 2021). Two additional modules (cyan and salmon) were strongly positively correlated with the green-yellow module (correlation coefficients 0.501 and 0.654, respectively) (Fig.S3). Although no pathways were identified in the salmon module, the cyan module showed a neurodegenerative signature (Table S2). Both cyan and salmon modules showed an enrichment of genes encoding mitochondrial proteins (16% and 10%, respectively) (Fig.2C). These data align with prior work that suggests there is a metabolic component to neurodegenerative disease (Muddapu et al., 2020; Pérez et al., 2018; Toledo et al., 2017). PGC-1a is the master regulator of nuclear encoded mitochondrial genes and is encoded by the Ppargc1a gene. RT-PCR using primers that detect all transcript isoforms derived from Pparc1a identified significantly lower levels in mice of advanced age compared to young and middle-aged adult mice (Fig.2D). These findings suggest that changes in mitochondrial functional pathways are a significant factor in brain aging, that metabolic and neurogenerative pathways are related to each other and coordinated in their response to aging, and these age-related changes are associated with lower expression of PGC-1a.

**Figure 2:**
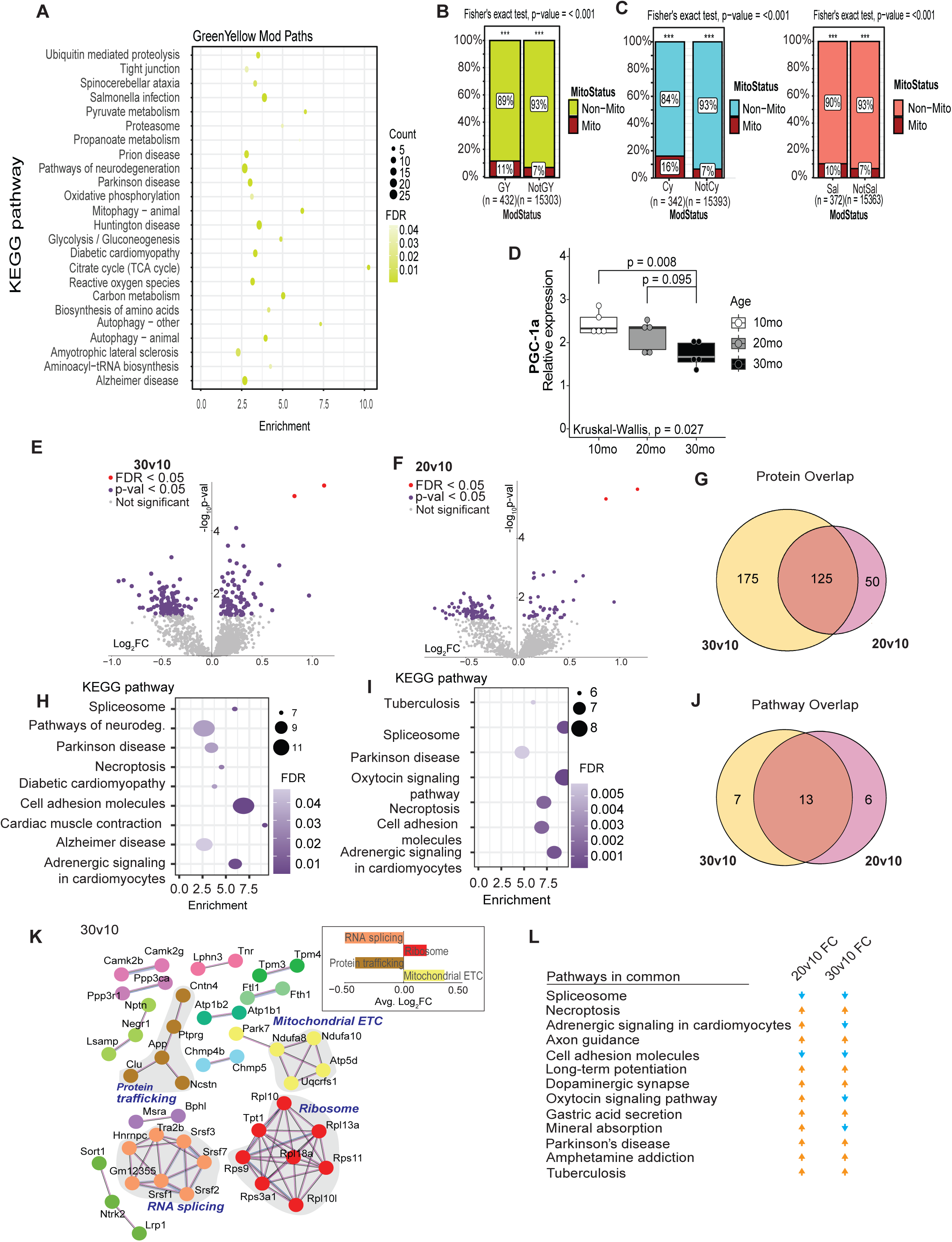
Metabolism of brain aging is reflected in the transcriptome and proteome. **A)** KEGG pathways detected in the green-yellow module of WGCNA from age male mouse cortex. **B-C)** Abundance of Mitocarta 3.0 genes in the green-yellow (B), cyan, and salmon (C) modules. **D)** RT-qPCR detection of PGC-1a expression cortex of male mice at 10 months, 20 months, and 30 months old (n=5). **E-F)** Volcano plots displaying proteins quantified. Statistically significant proteins are highlighted in purple. **G)** Venn diagram showing the overlap between proteins that were differentially expressed in the 30-month-old mice or 20-month-old mice compared to the 10-month-old. **H-I)** KEGG pathways enriched in the 30-month-old cortical proteome (H) and 20-month-old cortical proteome (I). Only pathways with greater than 6 proteins were plotted. **J)** Venn diagram showing the overlap between enriched KEGG pathways in the 30-month-old mice, or 20-month-old mice compared to the 10-month-old cortex. **K)** Network identified by String analysis. Proteins are colored based on biological pathway - mitochondrial ETC (yellow), ribosome (red), protein trafficking (brown), RNA splicing (orange). **L)** Table displaying pathways in common between the 20v10 and 30v10 comparison and if they are activated or repressed compared to the 10-month-old mice.

Although transcriptional changes provide useful information, changes at the transcript level do not necessarily predict changes at the proteome level, and the overlap between profiling platforms is often low (Maier et al., 2009; D. Wang et al., 2019). Proteins were isolated from the same brain hemispheres using the organic phase of the extracts that were used to generate the RNA-Seq data and analyzed by liquid chromatography-tandem mass spectrometry (LC-MS/MS). Across all samples, the detected proteome included 3578 quantified proteins, of which 2651 were annotated (Table S4). Comparison across age groups identified 300 proteins as different in abundance between 30-month-old mice and 10-month-old mice (unadjusted p < 0.05), and 175 that differed between 20-month-old and 10-month-old mice (Fig.2E; Fig.2F). Of those proteins that were responsive early during aging, ∼70% persisted through the later aging phase (Fig.2G). Functional enrichment of the protein differences for each comparison was assessed via overrepresentation analysis (ORA) using KEGG (Fig.2H; Fig.2I). Pathway overlap between age groups was 50% (Fig.2J), including those associated with neurodegeneration (Parkinson’s and Alzheimer’s, both of which include oxidative phosphorylation components), cell adhesion, adrenergic signaling, and RNA processing (spliceosome) (Fig.2I; Fig.2J). Taking an alternate approach, connections among age-sensitive pathways were also investigated via string (Fig.2K). Functional enrichment analysis identified mitochondrial electron transport chain (ETC) and ribosomal pathways that were positively enriched and protein trafficking and RNA splicing pathways that were significantly downregulated in response to age (Tukey’s, nominal p<0.05). Finally, changes from the early to later aging phases were largely congruent in terms of the directionality of change (Fig.2L), indicating that at least some of the age-associated changes in the brain at the protein level are progressive and are initiated between 10 and 20 months of age. Overall, proteomics analysis demonstrates that aging induced changes in mitochondrial pathways and processes linked to either growth or maintenance of neuronal function are coincident.

### PGC-1a transcript isoforms in the brain and cell type specificity

The brain is unique in producing variants of PGC-1a driven from a distal promoter located ∼590kb upstream from the canonical promoter of the Ppargc-1a gene (Fig.3A). The brain promoter contains up to five brain-specific exons spliced to canonical exon 2 of the PGC-1a gene (Soyal et al., 2012). All possible isoforms from the brain promoter can be detected simultaneously using primers against brain exon 1 and canonical exon 2 (Table S5), with the product hereafter referred to as PGC-1a B1E2. The isoforms generated from this transcript are predicted to encode proteins almost identical to that generated from the canonical promoter, PGC-1a (PGC-1a1), with small differences appearing only at the N terminus (Soyal, Felder, Auer, Hahne, Oberkofler, Witting, Paulmichl, Landwehrmeyer, Weydt, Patsch, et al., 2012). In cortices from 1-year old C3B6-F1 hybrid male mice, isoform specific primers detected PGC-1a B1E2 transcript at ∼8-fold greater abundance than canonical full-length PGC-1a1 and ∼4.5-fold at greater abundance than that driven from the alternative promoter, PGC-1a4 (n=6, p<0.0001) (Fig.3B). To understand how isoform expression in bulk tissue corresponded to cell types, primary neurons and primary astrocytes were isolated from postnatal mouse brains, cultured, RNA extracted, and PGC-1a isoforms were detected. Remarkably, neurons and astrocytes differed completely in PGC-1a expression profiles. Neurons displayed ∼8-fold greater levels of PGC-1a B1E2 than PGC-1a1, with an intermediate expression of PGC-1a4 (Fig.3C). PGC-1a B1E2 was not expressed in astrocytes; instead PGC-1a1 was the most abundant isoform, with a minor contribution from PGC-1a4 (Fig.3D). Isoform specific primers were used to probe mRNA extracted from brain hemispheres of mice aged 10 months, 20 months, or 30 months, and showed that PGC-1a1, PGC-1a4, and PGC-1a B1E2 all decrease with age (p<0.05 for all isoforms) (Fig.3D).

**Figure 3:**
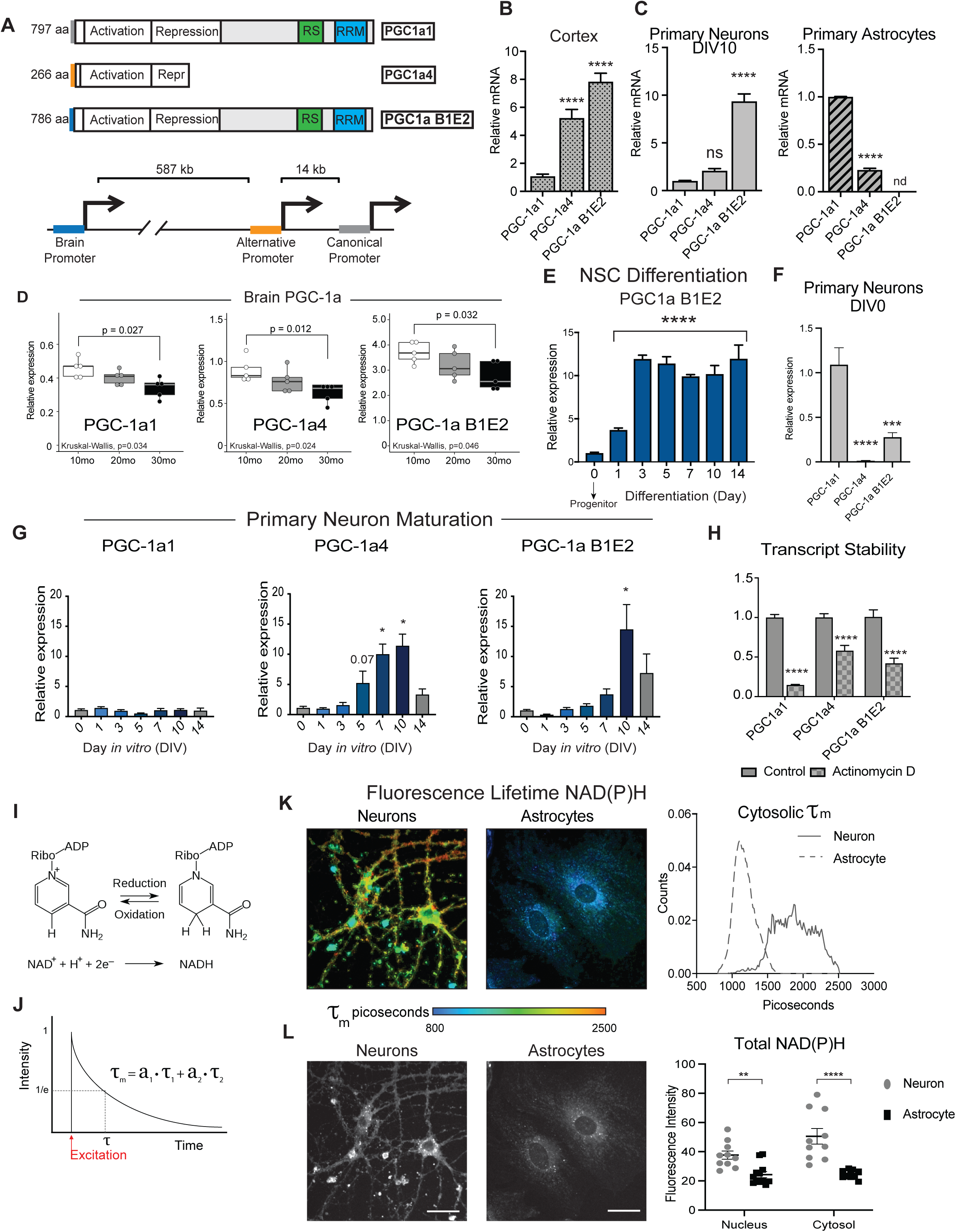
PGC-1a transcript isoforms in the brain and cell type specificity. **A)** Schematic of the three major isoforms of Ppargc1a expressed in the brain and a representation of the location of their three distinct promoter regions. **B-C)** Relative expression pattern of PGC-1a transcripts detected by RT-qPCR of 12-month-old male mouse cortex (n=6) (B) and DIV10 primary cortical neurons and astrocytes isolated from P0 (neurons) and P1 (astrocytes) neonates (n=6) (C). **D)** RT-qPCR detection of Ppargc1a transcript variants in cortex of male mice at 10 months, 20 months, and 30 months old (n=5). **E)** Expression of the brain-specific isoform of PGC-1a during neural stem cell (NSC) differentiation (n=3). **F)** Relative expression pattern of PGC-1a transcripts detected in DIV0 P0 neurons by RT-qPCR (n=6).**G)** Detection of Ppargc1a transcript variants during maturation of P0 primary cortical neurons (n=3-6). **H)** PGC-1a transcript levels after 24 hours of actinomycin D treatment (n=7) **I)** Schematic of oxidation-reduction reaction of NAD^+^ to NADH. **J)** Example of two component decay curve produced through fluorescence lifetime imaging microscopy and is represented by the equation τ_m_ = a_1_(τ_1_) + a_2_(τ_2_). **K)** Representative images (left) and distributions (right) of NAD(P)H mean fluorescence lifetime images of primary neurons and primary astrocytes (n=10-12). **L)** Representative images (left) and quantitation (right) of NAD(P)H fluorescence intensity of primary neurons and primary astrocytes (n=10-12). Statistical significance determined by ANOVA (B, C, D, E, F, and G) and student’s t-test (I). Asterisks indicate p value of <0.05 (*), <0.01(**), <0.0001 (****).

Neurons and astrocytes share a common progenitor; however, they differ in the expression of B1E2, suggesting that the brain promoter must be activated explicitly at some point during neuronal differentiation. To define when this happens, hippocampal neural stem cells (NSCs) were isolated, differentiated, and sampled at intervals over the 14-day differentiation process. PGC-1a B1E2 was barely detected in NSCs, increased ∼12-fold by day 3 of differentiation, and persisted at an elevated level for the remainder of the differentiation process (Fig.3E). In contrast, PGC-1a1 levels were reduced nearly 2-fold by day three and recovered by day 14, with a similar detected for PGC-1a4 over this same time frame (Fig.S4). Primary neurons isolated from neonate and prenatal brains require several days in culture to fully mature and to express classic neuronal transcriptional signatures. To understand the dynamics of PGC-1a isoform expression during primary neuron maturation, neurons were isolated from P0 mouse cortices and monitored over 14 days. On the initial day of isolation (DIV 0), PGC-1a4 was nearly undetectable, while PGC-1a B1E2 was 4-fold lower than PGC-1a1 (Fig.4F). During maturation, PGC-1a1 did not change, increased expression of PGC-1a4 was detected by day 5 of maturation, and increased expression of B1E2 at day 10 (Fig.3G). To determine if there were differences in stability of the neuronal transcript isoforms of PGC-1a, primary neurons were treated with transcription inhibitor actinomycin D. After 24 hours, PGC-1a1 levels were reduced by 85%, PGC-1a B1E2 was decreased by 58%, and PGC-1a4 was reduced by 43% (Fig.3H). The basis for differences in stability is unclear, in particular between PGC-1a and B1E2, which differ only in the 5’ region. These data suggest that neuronal PGC-1a transcript isoforms are not regulated equivalently or in parallel.

**Figure 4:**
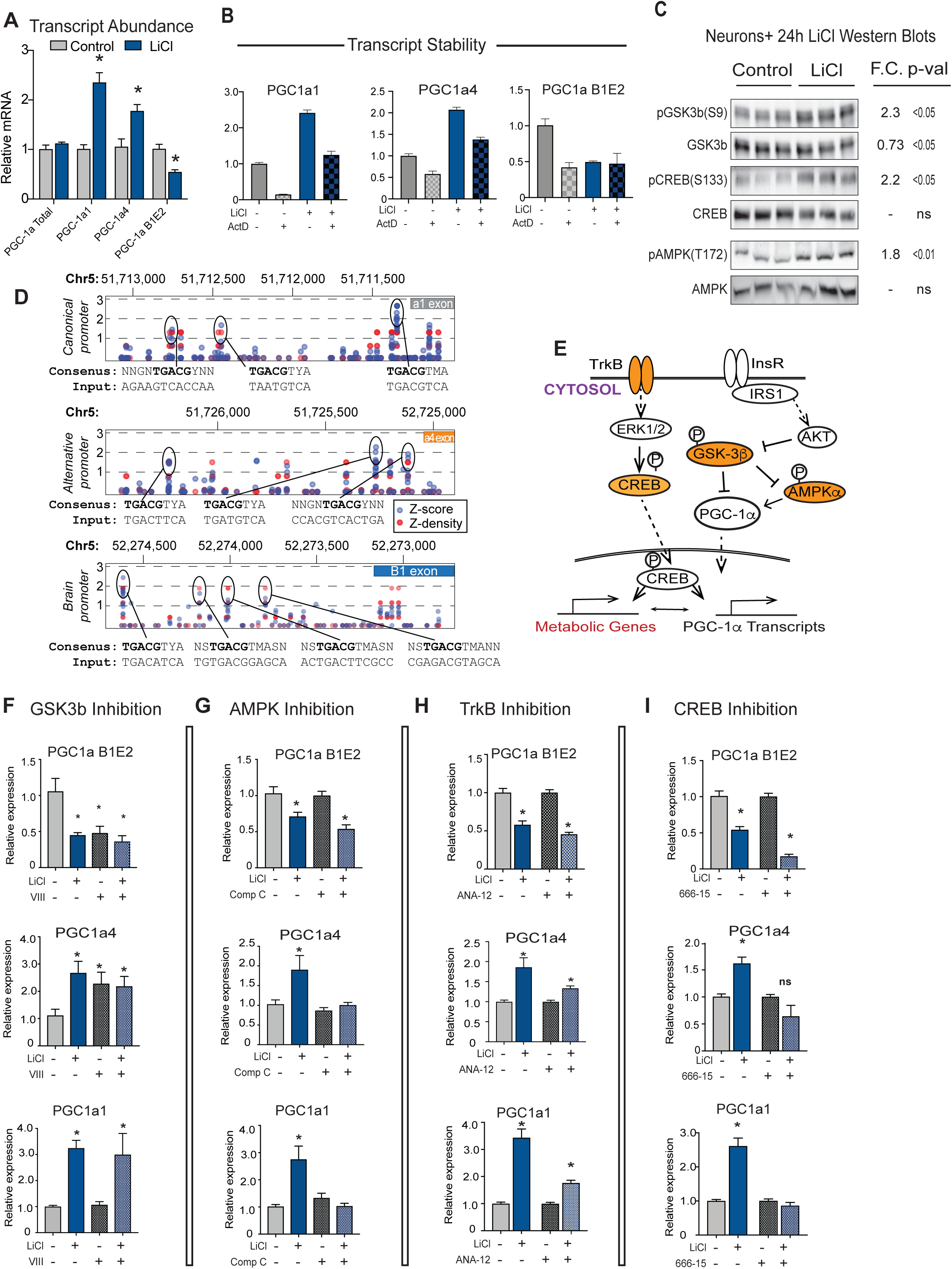
PGC-1a gene promoter activation by GSK3b and associated factors. **A)** RT-qPCR detection of PGC-1a transcripts in control and LiCl-treated neurons (n=6). **B)** RT-qPCR detection of PGC-1a transcripts after 24-hour treatment with actinomycin D +/-LiCl (n=6-7). **C)** Immunoblots of phosphorylation and total protein of GSK3β, CREB, and AMPK after 24-hour treatment with 15mM lithium chloride (LiCl) (n=6-7). **D)** Prediction of CREB binding sequences in Mmu Ppargc1a promoters using a transcription factor binding motif prediction software (http://tfbind.hgc.jp/). **E)** Schematic of proposed pathways of lithium regulation of Ppargc1a transcripts (orange indicates proteins investigated in F-I. **F-I)** RT-qPCR detection of Pparc-1a transcripts following treatment with lithium in the presence or absence of GSK3b inhibitor VIII (F), AMPK inhibitor Compound C (G), TrkB inhibitor ANA-12 (H), or CREB inhibitor 666-15 (I) (n=6-8). Asterisk (*) indicates p-value < 0.05 by student’s t-test (A-C) or 2-way ANOVA (F-I).

To understand if differences in PGC-1a isoform expression between primary neurons and primary astrocytes correlated with differences in cellular metabolic status, fixed cells were analyzed by fluorescence lifetime imaging microscopy (FLIM). This method allows for an independent assessment of the relative contribution of bound and free NAD(P)H to total NAD(P)H pools by fitting the decay curve to fast (t_1_) and slow (t_2_) photon release that are associated with free and bound fluorophores respectively (Fig.3I; Fig.3J). Mean fluorescence lifetime (*τ*_m_) was significantly higher in neurons than in astrocytes (Fig.3K). Analysis of lifetime components of NAD(P)H revealed that the neuronal decay curve (*τ*_m_) was comprised of a greater contribution from free than bound cofactors (70/30 ratio), but the decay curve in astrocytes had an even greater contribution (80/20) from free cofactor compared to bound (Fig.S5). These data suggest that there may be greater reliance on the TCA cycle and electron transport chain in neurons (protein-bound NADH) in primary neurons compared to astrocytes, an outcome that was not unexpected (Belanger et al., 2011). Total fluorescence intensity, a measure of cellular NAD(P)H, was higher in neurons than in astrocytes for both nuclear and cytosolic pools (Fig.3L). Taken together, these data show brain PGC-1a transcript expression changes with age across all three promoters, that PGC-1a B1E2 emerges through differentiation and maturation as neurons commit to their fate, and that cell-type differences in PGC-1a isoform expression between neurons and astrocytes are linked to distinct metabolic states.

### PGC-1a gene promoter activation by GSK3b and associated factors

In cultured cells, PGC-1a activity and protein stability can be modulated through the growth sensitive kinase GSK3b (R. M. Anderson et al., 2008; Martin et al., 2018). In mice, immunodetection of GSK3b expression in the hippocampus from the same 10, 20, and 30 months of age mice revealed significant age and region effects where GSK3b was increased at the later age phase (Fig.S6, 2-way ANOVA p<0.01). In glioma cells (H4) and in phaeochromocytoma-differentiated neurons (PC12), GSK3b inhibition using lithium chloride (LiCl) causes an increase in expression of PGC-1a1 and PGC-1a4 isoforms (Martin et al., 2018), but neither cell type expresses the B1E2 isoform which is the dominant form of PGC-1a in primary neurons. Neurons were treated with lithium chloride for 24 hours, and isoforms of PGC-1a were detected by RT-PCR. Unexpectedly, B1E2 expression from the brain promoter was suppressed by LiCl treatment even though, consistent with our prior reports, PGC-1a1 and PGC-1a4 transcripts were induced (Fig.4A). Changes in isoform expression were not observed at all in astrocytes which appeared completely refractory to lithium treatment (Fig.S7). These data suggest that mechanisms controlling the expression from the various neuronal promoters are not equivalent and differ by cell type.

To investigate potential differences in mRNA stability, neurons were treated for 24 hours with LiCl in the presence or absence of Actinomycin D. If mRNA stability were to explain the observed effect of LiCl then the expectation would be PGC-1a1 and PGC-1a4 stabilized and B1E2 de-stabilized, but this is not what was observed. LiCl modestly increased the stability of PGC-1a1 and B1E2 with no significant change in the truncated alternative isoform PGC-1a4 (Fig.4B). Next, a candidate approach was used to identify factors that might contribute to LiCl-directed regulation of Ppargc1a gene expression. Inhibitory phosphorylation of GSK3b (S9) was increased within 24 hours of treatment, as expected. LiCl also induced activating phosphorylation of CREB (S133), a transcription factor known to regulate PGC-1a expression (Finck, Brian N., and Kelly, Daniel P., 2006; Handschin et al., 2003), and AMPK (T172), another PGC-1a activator (Fig.4C). Prior studies had placed CREB downstream of GSK3b (Grimes & Jope, 2001), but a connection between AMPK and GSK3b has not been described. Analysis of canonical CREB binding motifs at each of the PGC-1a promoters revealed multiple putative CREB binding sites in each of the promoter regions of the Ppargc-1a gene (Fig.4D), arguing that LiCl may act in part through CREB. These and other factors were tested for their role in LiCl-directed regulation of PGC-1a (Fig.4E). LiCl is not a specific GSK3b inhibitor (Brown & Tracy, 2013), so to directly test for GSK3b regulation of PGC-1a neurons were treated with a specific inhibitor called inhibitor VIII. Treatment with inhibitor VIII phenocopied the effects of LiCl on PGC-1a4 (activated) and on B1E2 (repressed) but not on PGC-1a1 that only responded to LiCl and not inhibitor VIII (Fig.4F). Inhibition of CREB using inhibitor 666-15 had no impact on basal PGC-1a expression across all three isoforms but blunted the activation of PGC-1a1 and PGC-1a4 transcripts in response to LiCl, indicating that CREB is required. LiCl and CREB inhibition appeared to have an additive effect in repressing B1E2 (Fig.4G), suggesting that additional factors signaling to CREB are also involved. Treatment of neurons with an AMPK inhibitor, compound C, blocked the activation of PGC-1a1 and PGC-1a4, but the GSK3b-directed repression of B1E2 was not dependent on AMPK (Fig.4H). Going upstream in the signaling pathway, neurons were next treated with ANA-12, a TrkB inhibitor that blunts but does not entirely block the increase in expression in PGC-1a1 and PGC-1a4 and had no impact on B1E2 (Fig.4I). These experiments show that the canonical and alternative promoters are regulated in similar fashions, at least in response to LiCl, requiring TrkB, CREB, and AMPK for full implementation of the response to GSK3b inhibitors; however, the brain-specific promoter seems to be controlled by an alternate mechanism downstream GSK3b inhibition involving an as yet unidentified factor.

### Neuronal metabolic response to GSK3b inhibition with LiCl

The consequences of the repression of B1E2 PGC-1a isoform were investigated further via RNA-seq of LiCl treated neurons compared to untreated controls (n=4 per group) transcripts representing 15,370 genes were identified. Of those, 5,480 were identified as differentially expressed using an FDR of 0.01. Top LiCl responsive genes included ATF3, a cAMP-responsive transcription factor, solute carriers Aquaporin 9 and the zinc transporter Slc20a3 associated with synaptic vesicles, and homeobox transcription factors Pou4f3 and Dmbx1 (Table S6). Differentially expressed genes were subject to pathway analysis via GSEA (Fig.5A). Enriched and activated pathways included those associated with growth, such as cell cycle, Hippo signaling, and PI3K-Akt signaling that is part of the insulin and mTOR signaling pathway. Ordinarily, GSK3b is inhibited by growth signaling, but these data indicate that GSK3b inhibition as an initiating event might act to stimulate growth pathways, at least in neurons. The most highly enriched and suppressed pathways include ribosome, oxidative phosphorylation, proteosome, and several neurodegenerative disease pathways. Notably, many pathways that were repressed by LiCl were among those activated by aging in the brain, including oxidative phosphorylation (Fig.5B) and ribosome (Fig.5C). Focusing on established targets of PGC-1a transcriptional co-activation, a suite of genes associated with the electron transport chain (Cox4i1, Ndiufb8, Cycs, Vdac1, Cox5b), TCA cycle (Sdhb, Idh3a), and antioxidant maintenance (Sod2 and Cat) appear downregulated in response to LiCl, while expression of other PGC-1a associated factors including Nrf1, Ppara, Ucp2 were induced (Fig.5D). Genes associated with the neuronal function (Syt1 and Cplx1 (Lozoya et al., 2021) appear to be the most down-regulated, supporting PGC-1a B1E2’s role in modulating neuron-specific functions (Lozoya et al., 2021). A significant increase in BDNF expression, a neurotrophic factor important for neuronal health was also detected, and although BDNF is linked to PGC-1a levels and activation status at least in skeletal muscle (Wrann et al., 2013), these data suggest that the neuronal PGC-1a B1E2 isoform may be independent of BDNF.

**Figure 5:**
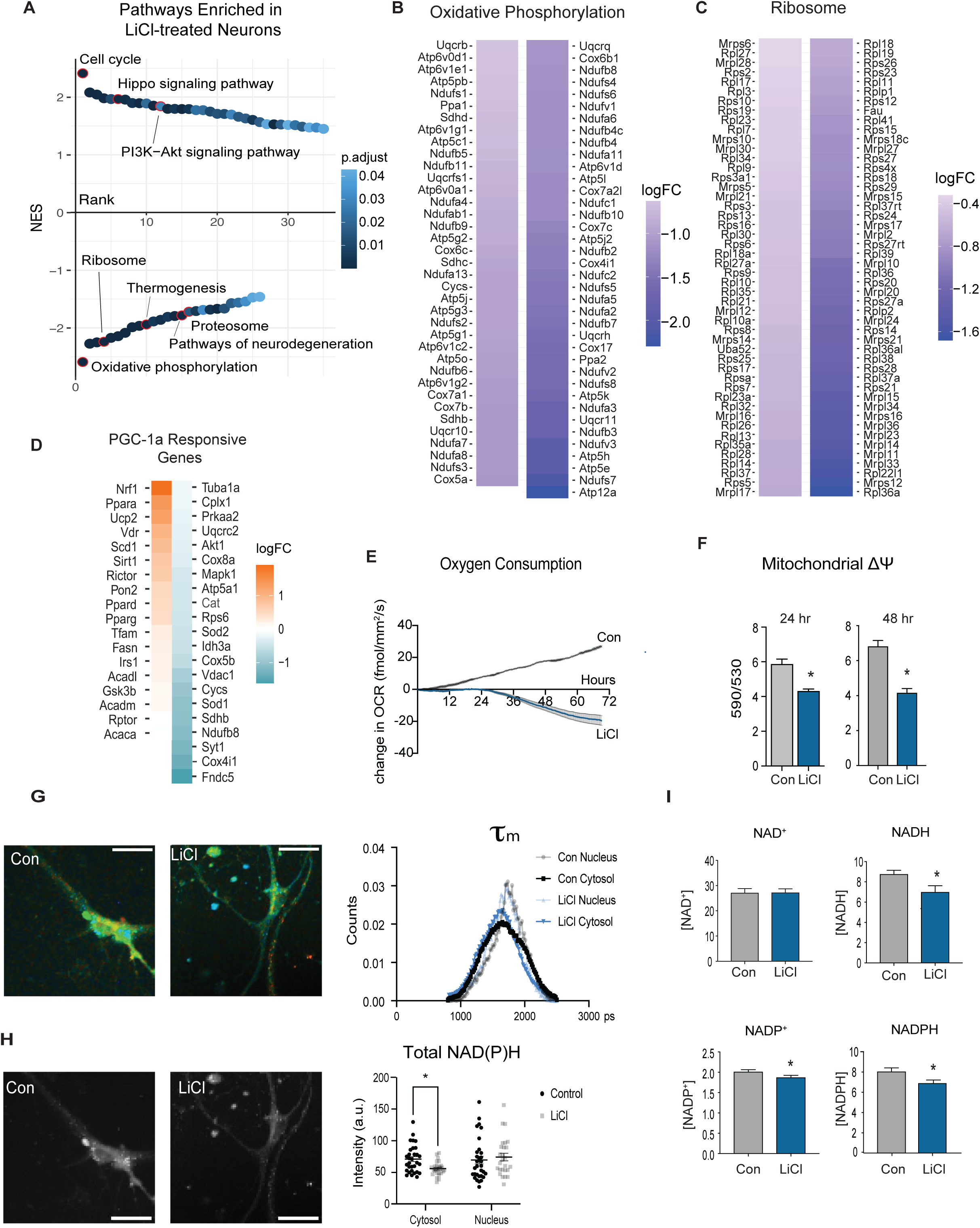
Neuronal metabolic response to GSK3b inhibition with LiCl. **A)** Rank order plot of enriched pathways detected by GSEA of RNA-sequencing of neurons treated for 24 hours with 15mM LiCl (n=4). **B-C)** Heatmap of the significantly changing genes in the oxidative phosphorylation (B) and ribosome (C) pathways enriched in LiCl-treated neurons. **D)** Heatmap of transcripts detected in the LiCl-treated neuron RNA-sequencing data set known to be responsive to changes in PGC-1a expression ranked by logFC. **E)** Oxygen consumption of primary neurons treated with LiCl or a control media change measured by RESIPHER live-cell oxygen consumption monitor for 72 hours (n=6). **F)** Mitochondrial membrane potential assessed by JC-1 assay in primary neurons treated for 24 and 48 hours with 15mM LiCl. **G)** Representative images (left)and distributions (right) of NAD(P)H mean fluorescence lifetime of control or LiCl-treated primary neurons. **H)** Representative images (left) and quantitation (right) of NAD(P)H fluorescence intensity of control or LiCl-treated primary neurons. **I)** NAD, NADH, NADP, and NADPH biochemical assays. Asterisks indicate p-value <0.05 by student’s t-test (F, H, and I) and multiple Mann-Whitney tests (E).

Changes in gene expression suggested that LiCl would blunt oxidative metabolism in neurons. Oxygen consumption was quantified continuously in primary neurons treated with LiCl using the RESIPHER oxygen consumption monitor. By 24 hours, the oxygen consumption of the neurons in culture was significantly lower than that of untreated neurons and remained lower out to 72 hours (Fig.5E). Mitochondrial membrane potential was also lowered by 24hr LiCl treatment and remained lower than that of untreated neurons up to 48 hours (Fig.5F). Fluorescence lifetime imaging of redox cofactors NAD(P)H using 2-photon microscopy detected a decrease in mean fluorescence lifetime (*τ*_m_) in primary neurons treated with LiCl with an increased contribution from free NAD(P)H pools, consistent with the cells having lower reliance on oxidative metabolism (Fig.5G). Additionally, a decrease in cytosolic NAD(P)H fluorescence intensity was detected in primary neurons treated with LiCl, consistent with the cells having lower oxidative metabolic activity (Fig.5H). Biochemical assessment of redox ratios for NAD and NADP in LiCl-treated whole cell extracts are consistent with the FLIM results. In enzyme-linked assays, significantly lower levels of the reduced form, NADH, were detected, but no change in the oxidized form, NAD+ (Figure 5G). For NADP, NADP+ and NADPH levels were both lower in LiCl-treated neurons (Figure 5H). These data point to a change in the redox state of cells treated with LiCl and connect redox metabolism to GSK3b. The metabolic impact of LiCl in primary neurons contrasts with that of other cell types that do not express PGC-1a B1E2 (Martin et al., 2018). In all cases, LiCl induces transcripts from the alternate and canonical promoters, but the repression of expression from the brain promoter is unique to neurons. The effect of LiCl on lower levels of mitochondrial-associated metabolic processes in neurons points to a functional dominance of PGC-1a B1E2 above other isoforms and suggests a means to confer cell-type specificity in the metabolic response to growth via regulation of PGC-1a.

### GSK3b plays an integrating role in growth regulation

Prior studies in immortalized glia (H4) and differentiated neurons (PC12) suggested that growth and metabolism networks might be interconnected via the GSK3b/PGC-1a axis (Martin et al., 2018). To determine whether these processes also intersect and interact in primary neurons, the role of GSK3b inhibition in controlling the growth signaling in neurons was investigated. Treatment with LiCl resulted in higher levels of inhibitory phosphorylation of insulin receptor substrate -1 (IRS-1) in the phosphatidyl inositol 3 kinase (PI3K) binding domain (S632) and lower levels of activating phosphorylation of AKT (T308) and S6 ribosomal protein (S240) (Fig.6A). These differences in post-translational modification are consistent with the repression of the insulin/IGF and mTOR Complex 1 signaling pathways. Additionally, LiCl induced higher levels of activating phosphorylation of ERK1/2, a key effector of BDNF-TrkB signaling and known kinase for CREB. These data, together with the activation of CREB and AMPK (Fig.4C), place GSK3b at the center of an interwoven growth and metabolism regulatory network (Fig.6B).

**Figure 6:**
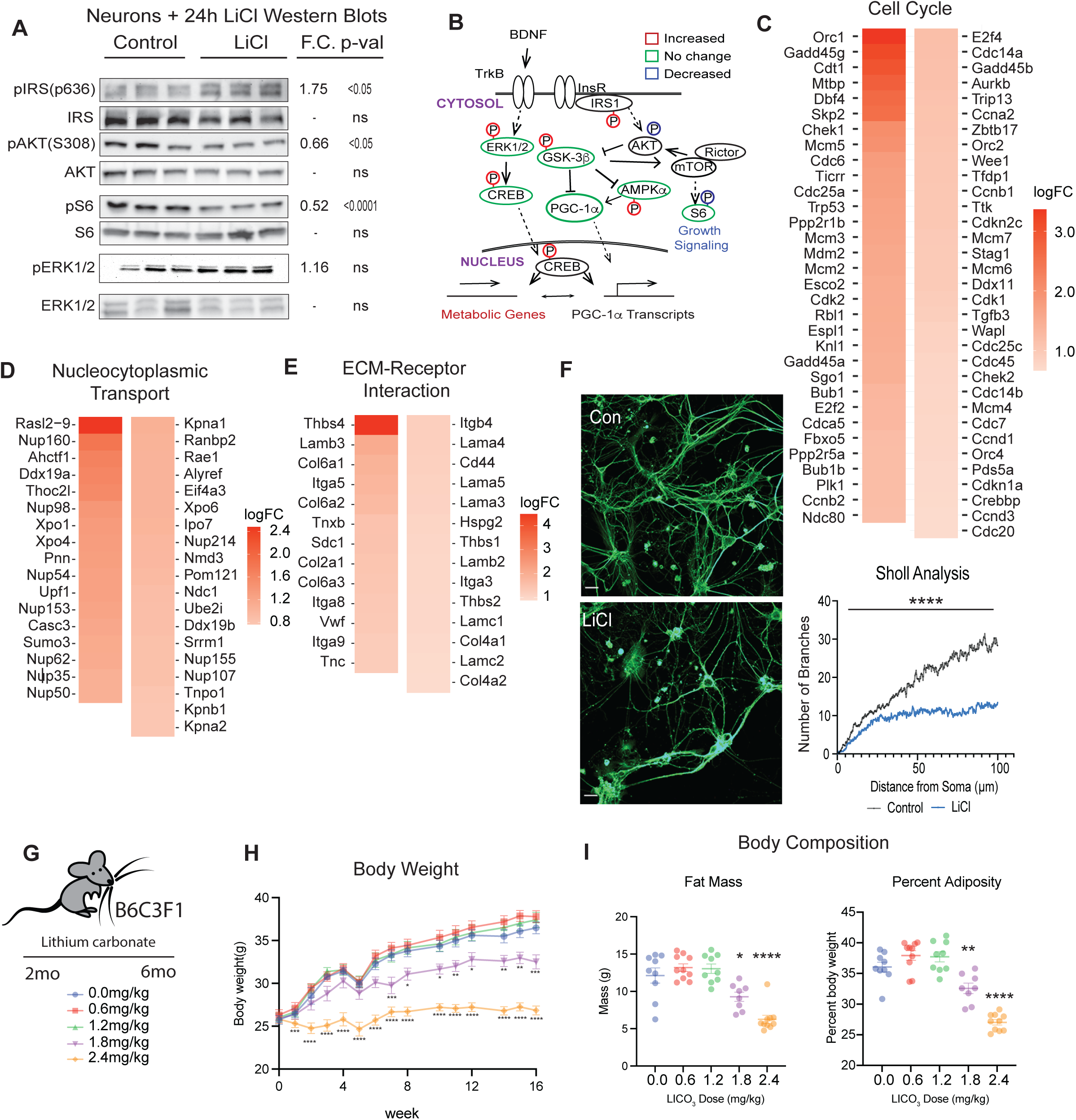
GSK3b plays an integrating role in growth regulation. **A)** Immunodetection of phosphorylated and total protein levels of IRS, AKT, S6, and ERK1/2 in primary neurons (n=6-8). Statistical significance was determined by Student’s t-test. **B)** Schematic of protein expression and phosphorylation in LiCl-treated neurons by Western blot (Figs. 4C and 5A) (n=6-8). **C-E)** Heatmaps of cell cycle, nucleocytoplasmic transport, and extracellular matrix-receptor interaction pathways detected GSEA in lithium-treated neuron RNA-sequencing. Heatmaps are ordered by log_2_FC. **F)** Representative images (left) and quantitation (right) of immunodetection of tubulin in control and LiCl-treated primary neurons (n=13-15). Quantitation was done by Sholl analysis using the Sholl Analysis plugin in ImageJ (**** indicates p<0.0001 determined by Student’s t-test at each distance). **G)** Schematic of feeding timeline for lithium carbonate mouse study. Mice were fed lithium carbonate-containing food at 0, 0.6, 1.2, 1.8, and 2.4 mg/kg daily for 16 weeks (n=8-10 mice). **H)** Body weights of the mice fed diets containing the five doses of LiCO_3_. Data shown as mean ± SEM. Statistical significance determined by ordinary 2-way ANOVA. **I)** Body composition analysis for the LiCO_3_ fed mice. Total fat mass (Left) and fat mass as a percent of total adiposity (right) are shown. Individual data is shown with error bars noting mean ± SEM. Statistical significance was determined by ordinary one-way ANOVA.

At the transcriptional level, Cell Cycle was the top-ranked enriched pathway (Fig.6C). At first glance, this might be an unexpected outcome since primary neurons are terminally differentiated; however, many of the genes within that pathway are also involved in cytoskeleton rearrangement, DNA surveillance, repair, and maintenance, and chromatin remodeling. Along these lines, the p53 pathway was ranked third and included several of the same genes (Table S5). The second-ranked pathway was Nucleo-cytosolic transport, including structural and transporting components, indicating that LiCl induces a nuclear response not limited to changes in gene expression (Fig.6D). Under microscopy, LiCl appeared to cause changes in neuronal morphology, an observation supported by the identification of the ECM-receptor pathway among those highly enriched (Fig.6E). Within this pathway, 9 of the top 10 have been linked to neuronal outgrowth, while another, syndecan 1 (Sdc1) is involved in neuronal progenitor maintenance and proliferation (Q. Wang et al., 2012). Immunodetection of tubulin networks in untreated and LiCl-treated primary neurons revealed significant changes in cell shape and in the length and in dendritic branching that were quantified using Sholl analysis (Fig.6F). These data show that GSK3b inhibition is associated with changes in intracellular growth signaling and in pathways related to growth, that this signature is transmitted to nuclear processes and has functional consequences for neuronal morphology and changes in neurite outgrowth.

To understand whether the impact of lithium on metabolism and growth might be conserved in vivo, C3B6F1 hybrid male mice were fed a diet supplemented with lithium for 4 months (Martin et al., 2018). Lithium carbonate (Li_2_CO_3_; 0.6, 1.2,1.8, or 2.4mg/kg) was incorporated into the diet of individually housed mice that were fed daily equivalent amounts of food starting at 2 months of age (Fig.6G). At the higher doses (1.8 and 2.4 mg/kg) mice gained less weight than their counterparts that were untreated or on lower doses (Fig.6H). Body composition analysis via DEXA (dual x-ray absorptiometry) showed that differences in body weight were explained entirely by differences in adiposity (Fig.6I). These data show that Li_2_CO_3_ treatment dampens growth at the whole organism level, favoring a reduction in adiposity without compromising lean mass, and indicate that the connection between metabolism and growth via GSK3b identified in cultured cells may also be relevant at the systems level.

### LiCl impacts brain growth and metabolism pathways

The effect of GSK3b inhibition on brain transcriptome was investigated via RNA-Seq using brain hemispheres from C3B6-F1 hybrid male mice (n=4 per group) treated with Li_2_CO_3_ (0.6, 1.2, 1.8, and 2.4 mg/kg in the diet) (Fig.6G). The impact of Li_2_CO_3_ was dose-dependent, with increasing numbers of responsive transcripts detected as the dose increased, with 15, 91, 176, and 358 responsive genes for each of the 4 doses, respectively (Fig.7A; Table S7). Pathway analysis via GSEA using an FDR cutoff of 0.1 identified few pathways for any dose. PathfindR is an alternate approach that identifies active subnetworks of altered genes before performing the enrichment analysis (Ulgen et al., 2019). A greater number of lithium-responsive pathways was identified with each step up in drug dose, with 61, 108, and 126 for doses 1.2, 1.8, and 2.4 mg/kg, respectively (Fig.7B; Table S7). There was only one pathway in the low-dose (0.6 mg/kg), axon guidance, and it was detected in the analysis of all of the other doses.

**Figure 7:**
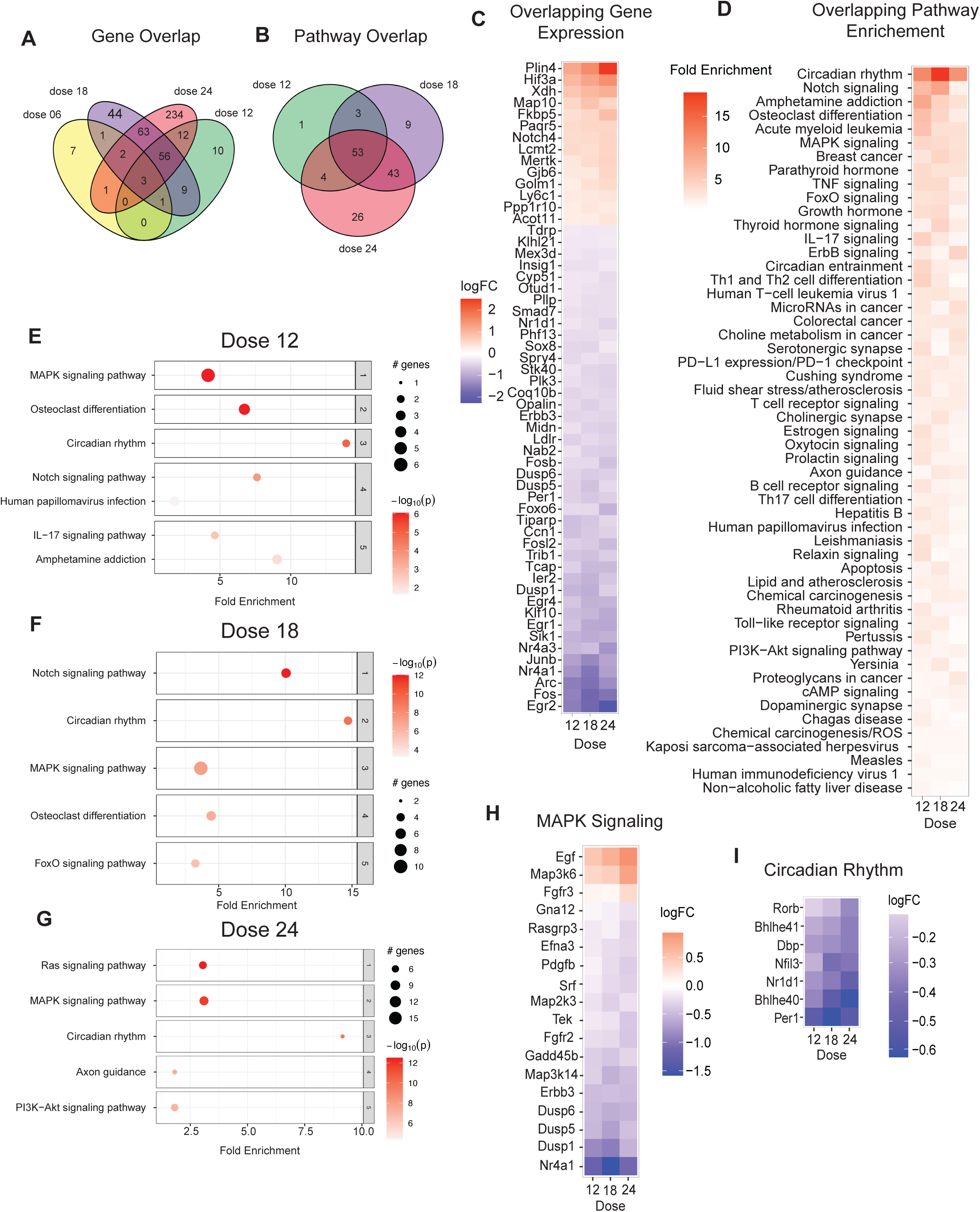
LiCl impacts brain growth and metabolism pathways. **A)** Venn diagram of significant genes detected in the brains of mice treated with 4 different doses of LiCO_3_ compared to control mice (n=4). **B)** Venn diagram of pathways enriched in the brains of mice given 1.2, 1.8, and 2.4 g/kg/day. Pathways were detected by PathfindR. **C)** Heatmap of the 56 genes that are differentially expressed in all 3 doses color coded by logFC. **D)** Heatmap of 53 overlapping enriched pathways detected in all three doses by PathfindR ranked by fold enrichment. **E-G)** Dotplots of top 5 enriched pathways in the 1.2 (E), 1.8 (F) and 2.4 (G) g/kg mouse brains. **H-I)** Heatmap of the significantly changing genes in MAPK Signaling (H) and Circadian Rhythm (I) pathways across each of 3 doses of LiCO_3_ 1.2, 1.8, and 2.4 g/kg.

Unlike the case with aging where genes were not uniformly changing across age groups, for this cohort, by and large, there was a progressive consistent change in levels of responsive genes across drug doses (Fig.7C). The overlapping genes among all doses included central transcription regulators responsive to growth, Fos and Junb, both of which were lower in the Li_2_CO_3_ treated animals. Several early growth response (EGR) genes were identified, including ERG2, ERG1, and ERG4, and all 3 were lower in abundance in Li2CO3-treated brains. Other genes of interest include Nr4a3, a target of p53, Sik, a serine-threonine kinase that regulates CREB, and Arc, a MAPK-responsive regulator of intracellular communication. Genes with higher expression included hypoxia-inducible factor 3a (Hif3a), a member of the HIF family lacking the transactivation domain that acts as a negative regulator of genes with HIF response elements. Plin4, a member of the perilipin family associated with lipid droplets, and Xdh, xanthine oxidase involved in purine degradation and salvage, both were higher in abundance with increasing doses of lithium carbonate.

Of the 53 overlapping pathways identified as being responsive to Li_2_CO_3_ treatment, the most highly represented signature related to immune and inflammatory pathways (34%), with similar representation from growth and metabolism pathways (46% combined) (Fig.7D). Two of the top 5 pathways (Amphetamine, Osteoclast) are largely explained by the presence of Fos, Jun, and Arc. Notch signaling, a key factor in juxtacrine communication, was ranked second among the Li_2_CO_3_ response pathways and included Notch4 and the notch pathway-associated transcription factor effector Maml3 that were increased in abundance, and Hes repressor proteins that were decreased in abundance (Table S7). The osteoclast differentiation pathway was also enriched and included Zbtb16 transcriptional repressor linked to HDACs and Rpskb2 that were increased in abundance, and Dusp6, a negative regulator of MAPK signaling, and Rara (retinoic acid receptor) involved in transcriptional regulation that was lower in abundance. Analyzing each dose individually, the relative enrichment of pathways in cortex from mice on each of the 1.2mg/kg (Fig.7E), 1.8mg/kg (Fig.7F), and 2.4 mg/kg (Fig.7G) doses was not equivalent (Fig.7E). At higher doses of Li_2_CO_3_ pathways linked to insulin signaling are identified including PI3k-Akt signaling (1.8mg/kg) and FoxO signaling (2.4mg/kg). The MAPK growth signaling (Fig.7H) and circadian rhythm (Fig.7I) pathways stand out as shared features when the data were analyzed in individual sets, and in general, the expression of genes in both pathways was downregulated by Li_2_CO_3_ in a dose-dependent manner.

## Discussion

Brain aging has long been associated with changes in inflammatory and neuronal functional pathways (Aron et al., 2022), but a more recently appreciated adaptation to age centers on metabolism (Camandola & Mattson, 2017; Cunnane et al., 2020). Conditions associated with aging in the brain include a greater risk for neurodegenerative disease and cognitive decline, both of which have been linked to disruption in mitochondrial processes (López-Doménech & Kittler, 2023; Trigo et al., 2022). GSK3b appears to be well poised at the center of a metabolism and growth regulatory network (Souder & Anderson, 2019), and this work suggests that its importance in brain aging not only relates to its impact on growth signaling pathways but also on its impact on metabolism via PGC-1a. In this study, transcriptional profiles of the cortex from male mice confirm the immune/inflammation and neuronal structural and communication pathways as the most responsive to age and reveal an expansive network of metabolic genes that are age-sensitive. This study agrees with the older literature in that changes with age are rather limited in terms of total numbers of differentially expressed genes (Lee et al., 2000); however, it is important to realize that aging introduces heterogeneity among individuals that can present challenges for meeting statistical significance. With advances in transcriptional and proteomic profiling, it has become clear that changes in the transcriptome are often not reflective of changes in the proteome (Maier et al., 2009). Obviously, there are profound differences in the range and sensitivity of detection between these two platforms, but another consideration is the importance of translational and post-translational regulation. Proteomic analysis confirms age-related changes in metabolism and processes vital for neuronal function but further shows changes in ribosomal and RNA splicing pathways. It is as yet unclear how changes in ribosomal pathways ultimately impact translation, including which transcripts are preferred and how selectivity might be altered. Changes in RNA processing pathways have been reported as a function of age and have recently been proposed as a potential hallmark of aging (Bhadra et al., 2020), although the significance of these alternations in terms of function remains unknown at this stage.

Although a role for mitochondria in aging is generally accepted, the role of “master regulator” PGC-1a in aging is not well defined. Despite its initial promise as a target for correcting age-related declines in metabolism (R. Anderson & Prolla, 2009), the issues of tissue-type and cell-type specificity in its expression, its interacting partners, and its target genes present a considerable challenge (Miller, Clark, & Anderson, 2019). In cultured cells, chronic low-level overexpression of PGC-1a impacts a host of cellular functions, including energetic, growth, and structural pathways, and these same pathways were found to correlate with PGC-1a expression in liver, adipose, and skeletal muscle from a genetically diverse cohort of mice (Miller, Clark, Martin, et al., 2019). In primary neurons, three different PGC-1a promoters are active and appear to be responsive to different regulatory inputs. At this stage, we can confirm that all 3 isoforms of PGC-1a are sensitive to changes in GSK3b activity and that CREB is important at all 3 promoters; however, the specifics of the influence of regulatory inputs such as GSK3b, AMPK, and CREB are distinct at each promoter. Neuronal metabolic status aligns with expression from the brain-specific promoter, indicating a functional dominance of this isoform. Genetic studies of PGC-1a B1E2 indicate that it regulates a suite of neuron-specific genes (e.g., neurotransmitter pathways) (Lozoya et al., 2021; Ruas et al., 2012), expanding its sphere of influence beyond energetics specifically in this cell type. In mice, PGC-1a was previously linked to arborization in neurons of the hippocampus (Cheng et al., 2012) and maintenance of the neuromuscular junction in skeletal muscle (Handschin et al., 2007), possibly secondary to its effects on mitochondria. Direct activities of PGC-1a include regulation of splicing (Martinez-Redondo et al., 2016) and nuclear to cytosolic mRNA transport (Mihaylov et al., 2023), each involving a mechanism that requires its RNA binding domain, present on PGC-1a B1E2 and on PGC-1a1 but absent on PGC-1a4. Neither of these activities have been specifically linked to PGC-1a B1E2, and the contribution of these regulatory mechanisms to neuronal function is as yet unknown.

The mood-stabilizing drug lithium has been well documented as a regulator of GSK3b and is thought to be effective through its influence on a variety of growth regulatory pathways, including Wnt, insulin/IGF, MAPK, and mTOR signaling (Patel & Woodgett, 2017). In mouse brains, GSK3b protein abundance increases with age, and in general, its expression distribution in the hippocampus and in the cortex mirrors that of PGC-1a. One of the unexpected outcomes of this study was the impact of GSK3b inhibition on primary neurons and astrocytes. The repression of PGC-1a at the brain promoter in response to LiCl or GSK3b inhibitor VIII is opposite to what is observed at the canonical and alternate promoters in neurons and in other cell types where the brain promoter is not activated, while the primary astrocytes were not at all responsive. This raises the possibility that, even in the absence of a GSK3b challenge, PGC-1a is part of a mechanism that allows different types of brain-resident cells to independently modulate cellular energetics and tailor their metabolic response to extracellular stimuli. The importance of GSK3b as a regulatory effector in central metabolism and growth signaling is underscored by the primary neuron and in vivo brain response to LiCl (neurons) and Li_2_CO_3_ (mice). It will be important to dissect how the neuronal PGC-1a response to GSK3b might be influenced by juxtacrine effects, in particular, whether contributions from astrocytes and microglia might modify the response reported here for neurons in primary culture.

A primary limitation of this study is that both of the in vivo studies, brain aging and the response to Li_2_CO_3_, were conducted in male mice only. Growing evidence points to profound sex dimorphism at the cellular and molecular level (Sampathkumar et al., 2020). Even while the physiological traits of aging manifest quite similarly for both sexes, it would appear that the key pathways and processes contributing to aging are quite sex dimorphic. For example, in rats, age-related changes in brain inflammatory cytokines have been reported for both males and females but appear to be more pronounced in females compared to males (Porcher et al., 2021). It will be important to uncover shared and sex-specific effects of aging on the brain and whether the interventions geared toward delaying aging might need to be optimized for sex. Despite this limitation, it seems clear that there are brain-specific mechanisms for controlling energetics and that changes in mitochondrial pathways are at least coincident with changes in inflammatory and neuronal functional pathways with age. Differences in how and when PGC-1a is activated are likely to play a role in establishing cell-type metabolic specificity, and differences in the PGC-1a response would allow for intracellular coordination in metabolic adaptation, where a single stimulus evokes distinct outcomes according to cell type. Given the breadth of pathways and processes that are influenced by mitochondrial status, there is considerable interest in advancing what we know about the contribution of mitochondria to disease and motivation to go beyond the concept of “dysfunction” (Monzel et al., 2023). Data from this study supports the idea that mitochondrial processes could be targeted to delay age-related changes in the brain and even potentially applied as a preventative measure for neurodegenerative disease.

## Methods

### Animals

#### Aging cohort

Six-week-old male B6C3F1 hybrid mice were obtained from Harlan Laboratories (Madison, WI, USA) and housed under controlled pathogen-free conditions in accordance with the recommendations of the University of Wisconsin Institutional Animal Care and Use Committee. Mice were fed 87 kcal week^-1^ of the control diet (Bio-Serv diet #F05312) from 2 months of age and were individually housed. This level of calorie intake is ∼95% of ad libitum for the B6C3F1 strain, so all food was consumed. By 30 months of age, the mortality of the control animals was ∼45%, consistent with the expected lifespan for this strain. Mice were euthanized by cervical dislocation at 10, 20, and 30 months of age.

#### Lithium diet study

Six-week-old male B6C3F1 hybrid were obtained from Harlan Laboratories (Madison, WI, USA) and housed under controlled pathogen-free conditions in accordance with the recommendations of the University of Wisconsin Institutional Animal Care and Use Committee. Mice were fed 87 kcal week^−1^ of the control diet (F05312; Bio-Serv, Flemington, NJ, USA) and were individually housed with ad libitum access to water. This level of food intake is ∼95% ad libitum for the B6C3F1 strain, so all food was consumed. Following two weeks of facility acclimation, mice were randomized into five treatment groups fed the control diet supplemented with increasing concentrations of dietary lithium carbonate (2 months old; n = 10/group): Group 1) 0.0 g/kg/day Li_2_CO_3_; Group 2) 0.6 g/kg/day Li_2_CO_3_; Group 3) 1.2 g/kg/day Li_2_CO_3_; Group 4) 1.8 g/kg/day Li_2_CO_3_; Group 5) 2.4 g/kg/day Li_2_CO_3._ Li2CO3-supplemented mice were administered an additional drinking bottle containing saline (0.45% NaCl) to offset polyuria, a common side effect of lithium treatment. Mice consumed dietary lithium for 4 months and were euthanized at 6 months of age. Mice were weighed every 1-2 weeks throughout the duration of the study. Body composition analysis was conducted using the Lunar PIXImus machine and software.

Brains were isolated, bisected, embedded in OCT, frozen in liquid nitrogen, and stored at −80°C until further processing.

#### Primary cell culture studies

Wild-type male and female C57BL6J mice were obtained from Jackson Laboratories (Bar Harbor, ME) and housed under controlled pathogen-free conditions in accordance with the recommendations of the University of Wisconsin Animal Care and Use Committee. Mice were allocated to breeding pairs at 8 weeks of age with two animals per cage.

#### RNA sequencing

RNA extraction was completed using a Direct-zol RNA kit (Zymo Research, Irvine, CA) according to the manufacturer’s instructions. Each RNA library was generated following the Illumina TruSeq RNA Sample Preparation Guide and the Illumina TruSeq RNA Sample Preparation. Purified total RNA was used to generate mRNA libraries using NEBNext Poly(A) mRNA Magnetic Isolation Module and NEBNext Ultra RNA Library Prep kit for Illumina (Illumina Inc., San Diego, CA, USA). Quality and quantity were assessed using an Agilent DNA1000 series chip assay and Invitrogen Qubit HS Kit (Invitrogen), respectively. Sequencing reads were trimmed to remove sequencing adaptors and low-quality bases (Jiang et al., 2014), aligned to mm10 reference genome using the STAR aligner, and alignments used as input to RSEM for quantification. Differential gene expression analysis was performed via EdgeR generalized linear model (GLM) method.

#### Proteomics

Flash-frozen mouse half brain samples were homogenized in digestion buffer [8M Urea with Proteinase inhibitors] then methanol:chloroform extracted per customer protocol. Protein pellets were resolubilized in 8M Urea without proteinase inhibitors. 100ug total protein per each of the 15 samples was reduced by incubation with 25mM DTT for 15 minutes at 56°C. After cooling on ice to room temperature 6μl of 55mM CAA (chloroacetamide) was added for alkylation and samples were incubated in darkness at room temperature for 15 minutes. This reaction was quenched by 16μl addition of 25mM DTT. Subsequent digestion was performed with Trypsin/LysC solution [100ng/μl 1:1 Trypsin (Promega):LysC (FujiFilm) mix in 25mM NH_4_HCO_3_] along with 40μl of 25mM NH_4_HCO_3_ (pH8.5) for a ∼1:40 substrate-to-enzyme ratio. Digests were carried out overnight at 37°C then subsequently terminated by acidification with 2.5% TFA [Trifluoroacetic Acid] to 0.3% final. 10% HFBA [Heptafluorobutyric acid] was added to 0.1% final, and each individual sample was cleaned up using OMIX C18 SPE cartridges (Agilent) according to manufacturer protocol. Eluates in 50%:50%:0.1% acetonitrile:water:TFA acid (vol:vol) were dried to completion in the speed-vac and reconstituted in 12ul of 0.1% formic acid and 3% acetonitrile. Peptides were analyzed by Orbitrap Fusion™ Lumos™ Tribrid™ platform, where 2ul of total proteome was injected using Dionex UltiMate™3000 RSLCnano delivery system (ThermoFisher Scientific) equipped with an EASY-Spray™ electrospray source (held at constant 50°C). Chromatography of peptides prior to mass spectral analysis was accomplished using capillary emitter column (PepMap® C18, 2µM, 100Å, 500 x 0.075mm, Thermo Fisher Scientific). NanoHPLC system delivered solvents A: 0.1% (v/v) formic acid, and B: 80% (v/v) acetonitrile, 0.1% (v/v) formic acid at 0.30 µL/min to load the peptides at 2% (v/v) B, followed by quick 2 minute gradient to 5% (v/v) B and gradual analytical gradient from 5% (v/v) B to 62.5% (v/v) B over 33 minutes when it concluded with rapid 10 minute ramp to 95% (v/v) B for a 9 minute flash-out. As peptides eluted from the HPLC-column/electrospray source survey MS scans were acquired in the Orbitrap with a resolution of 60,000, max inject time of 50ms and AGC target of 1,000,000 followed by HCD-type MS2 fragmentation into Orbitrap (36% collision energy and 30,000 resolution) with 0.7 m/z isolation window in the quadrupole under ddMSnScan 1 second cycle time mode with peptides detected in the MS1 scan from 400 to 1400 m/z with max inject time of 54ms and AGC target of 125,000; redundancy was limited by dynamic exclusion and MIPS filter mode ON.

Raw data was directly imported into Proteome Discoverer 2.5.0.400 where protein identifications and quantitative reporting was generated. Seaquest HT search engine platform was used to interrogate Uniprot Mus musculus (Mouse) proteome database (UP000000589, 63,686 total entries) along with a cRAP common lab contaminant database (116 total entries). Peptide cysteine carbamidomethylation was selected as static modifications whereas methionine oxidation and asparagine/glutamine deamidation were selected as dynamic modifications. Peptide mass tolerances were set at 10ppm for MS1 and 0.03Da for MS2. Peptide and protein identifications were accepted under strict 1% FDR cut offs with high confidence XCorr thresholds of 1.9 for z=2 and 2.3 for z=3. For the total protein quantification processing, label-free settings were used on unique and razor peptides, and protein grouping was considered for uniqueness. ANOVA (individual proteins) hypothesis was executed without imputation mode being executed.

#### Histochemistry

Serial cryostat sections of 10*μ*M in thickness were cut at -20*°*C with a Leica Cryostat (Fisher Supply, Waltham, MA, USA), defrosted and air-dried, and stained for GSK3*β* (9315; Cell Signaling Technologies, Danvers, MA, USA) as previously described(Pugh et al., 2013). Images were acquired on a Leica DM4000B microscope equipped with an HCX PL FLUOTAR 20x/ 0.50 objective. Photographs were taken with a Retiga 4000R digital camera (QImaging Systems, Surrey, BC, Canada). Camera settings were optimized for each stain, and for uniformity, all images were taken with identical settings, fixed light levels, and fixed shutter speeds.

### Cell culture

#### Primary Neurons

Primary cortical neurons were isolated from brains of P0 mice. Briefly, the cortices were harvested from pups into ice-cold HBSS where the midbrain and meninges were removed. Brain tissue was minced and digested in 0.25% trypsin for 20 minutes at 37*°*C. Trypsin was quenched with DMEM/10%FBS/1%Pen/Strep and cells were dissociated and counted prior to plating on poly-d-lysine coated plates. Cultures were then fully changed to Neurobasal Plus Media (2% B27^+^, 1% GlutaMax, 1% Pen/Strep) the subsequent day. Upon first change to Neurobasal media, neurons were treated with 1*μ*M AraC (cytosine arabinoside) to eliminate glial cell populations. Media was changed by ½ volume every 3-4 days until experiments. All primary neuron experiments were conducted between day 10-14 in vitro.

#### Primary Astrocytes

Primary cortical astrocytes were dissociated similarly to neurons and plated in DMEM/10% FBS/1% Pen/Strep. Briefly, cortices were collected from P1-2 mouse pups and pooled in groups of 2. Cells were dissociated enzymatically in 0.25% Trypsin for 20 minutes. Cells were further dissociated by trituration. Cells from 2 cortices were plated on a T-75 cell culture flask and grown in DMEM supplemented with 10% FBS and 1% Pen/Strep. Astrocytes proliferated to confluency and were subsequently treated with 10*μ*M AraC for 48 hours to eliminate non-astrocyte glial populations. Upon microglial removal, astrocytes were then passaged to a T-175 to ensure astrocytic growth. Cells were passaged for experiments at ∼80% confluence and plated for a one-day growth.

#### Neural stem cells (NSCs)

Hippocampi were dissected in cold HBSS and dissociated using the GentleMACS Dissociator (Miltenyi Biotec) and MACS Neural Tissue Papain Dissociation Kit (Miltenyi Biotec) according to the manufacturer’s protocol. NSCs were cultured in DMEM/F12 media (Invitrogen) supplemented with GlutaMAX, FGF (20ng/mL), EGF (20ng/mL) and Heparain (5ug/mL) for the indicated number of days prior to assay.

#### Drug Treatments

Cells were treated with 15mM LiCl 24h prior to metabolic assays, qPCR, and RNA-sequencing. For neurons, LiCl treatment accompanied a 50% media change. Drug treatments were co-administered with lithium during 50% media change as follows: Inhibitor VIII (Millipore) [15uM], CREB inhibitor 666-15 (Millipore) [1uM], ANA12 (Sigma-Aldrich) [20uM], Compound C (Millipore) [8uM].

#### RT-qPCR

Cells were treated as outlined above and RNA was collected 24 hours after treatment. For NSC differentiation and primary neuron maturation, cells were lysed at day in vitro indicated above. Cells were lysed with Trizol and RNA was isolated using Zymo Research Direct-zol RNA MiniPrep Kit. RT-qPCR was conducted using iTaq Universal SYBR Green Supermix (1725121, Bio-Rad, Hercules, CA, 94547). Primer sequences for all transcripts can be found in Table S5.

#### Multiphoton laser scanning microscopy

Conducted as previously described. Briefly, cells were grown on glass coverslips, fixed in 10% formalin, and mounted using Fluoromount (Thermo Scientific). Images were captured using a Nikon 60X 1.3apo objective and a 457/40 filter. The acquisition time for neuron-astrocyte comparison was 120 seconds. Lithium-treated neuron experiments were conducted using a 60-second acquisition time. Image analysis was conducted using SPCImage 8.0. (https://www.becker-hickl.com/products/spcimage/).

#### Promoter region transcription factor binding site determination

Predicted CREB binding sequences present in the promoter regions of the Ppargc-1a gene were predicted using transcription factor binding motif prediction software (http://tfbind.hgc.jp/). The prediction was made using the reported CREB-ChIP sequencing peaks identified by Lesiak et al. The strength of CREB prediction was based on the density score produced by the number of potential CREB binding sites. Confidence values from Lesiak et al., and density scores were then z-scored to determine the predicted binding sites.

#### Western Blot, and Immunofluorescence

Cells were lysed, and protein was extracted in a modified RIPA buffer containing protease and phosphatase inhibitors (P8340 and 524624, respectively; Sigma-Aldrich, St. Louis, MO, USA). Proteins were detected by immunoblotting using standard techniques. Antibodies used were all acquired from Cell Signaling Technologies unless otherwise noted and used at the manufacturer’s recommended dilution. Antibodies used were: pGSK3b(S9) (#9336), GSK3b (9315S), pCREB (S133) (ab32096; Abcam), CREB (#9104), pAMPK (T172) (#2535S), AMPK*α* (#2532S), pIRS (p636) (#2388S), IRS-1 (#2382S), pAKT (T308) (#13038S), AKT (#4691), pS6 (S240/244) (#2215S), S6 (#2217S), pERK1/2 (T202/ Y204) (#4370S), ERK1/2 (#4695S). Western blot densitometry was conducted in Adobe Photoshop to detect band intensity.

Immunofluorescence images were acquired on a Leica DM4000B microscope equipped with a Leica N Plan 40x/0.65 objective. Photographs were taken with a Retiga 4000R digital camera (QImaging Systems, Surrey, BC, Canada). Camera settings were optimized for each stain, and for uniformity, all images were taken with identical settings, fixed light levels, and fixed shutter speeds. Cells were grown on glass coverslips coated with poly-d-lysine. Cells were fixed in 10% formalin for 10 minutes, permeabilized with 0.3% Triton X-100 in PBS (PBS-T) for 1 hour, and incubated with anti-a-Tubulin primary antibody (1:200) (Sigma-Aldrich #T6199) diluted in 1% BSA overnight. Cells were washed with PBS and then incubated with a secondary antibody, Goat anti-mouse IgG,AlexaFluor 488, for 1 hour at room temperature. Coverslips were mounted using Fluoromount (Thermo Scientific). Sholl analysis was conducted with ImageJ (NIH, Wayne Rasband, http://rsb.info.nih.gov/ij/). using the Sholl Analysis plugin (https://imagej.net/plugins/sholl-analysis).

### Bioassays

#### Oxygen CONSUMPTION (OC)

OC was measured using a Resipher oxygen consumption monitor. OC was measured for 24 hours prior to treatment to establish basal respiration. Change in OC oxygen consumption rate was determined by comparing the oxygen consumption rate at 15-minute intervals taken for 24 hours post-treatment.

#### JC-1 Assay

Mitochondrial membrane potential was quantified with JC-1 dye (Invitrogen, Waltham, MA). Neurons were treated with 15 mM LiCl for 24 hours and then incubated with 1ug/mL JC-1 dye for 30 minutes. Cells were washed with PBS prior to fluorescence detection. Fluorescence was measured using excitation/emission wavelengths of 535/590 nm and 485/530 nm.

#### NAD(P) and NAD(P)H

NAD(P)+ and NAD(P)H concentrations were quantified with NAD(P)- and NAD(P)H-Glo assay kits (Promega) according to manufacturer instructions. Neurons were treated with 15 mM LiCl for 24 hours prior to assay.

#### Statistics

All student’s t-tests were two-tailed. Outliers were identified by Grubb’s test using a threshold of p < 0.05. One-way and two-way ANOVA were conducted assuming Gaussian distribution and corrected for multiple comparisons using Tukey’s test.

#### Contact for Reagent and Resource Sharing

Further information and requests for resources and reagents should be directed to Lead Contact Dr. Rozalyn Anderson.

## Acknowledgments

This work was supported by NIH R01AG067330, NIH training fellowships AG000213 (DCS), T32DK007665(ERM). This study was conducted using resources and facilities at the William S. Middleton Memorial Veterans Hospital, Madison, WI. The authors declare no conflict of interest.

## Conflict of Interest

The authors declare no conflict of interest.

## Notes

### Competing Interest Statement

The authors have declared no competing interest.

