## Supplemental Figure Legends for "Mitochondrial regulator PGC-1a in neuronal metabolism and brain aging"

Figure S1: **A)** Volcano plots displaying transcripts quantified. Statistically significant transcripts are highlighted in red (upregulated) or purple (downregulated) in 30-month-old compared to 10-month-old male cortical tissue as shown in figure 1A. Equivalent volcano plots for 30-month-old compared to 20-month-old and 20-month-old compared to 10-month-old are shown. **B)** Rank order plot of enriched pathways detected by GSEA. Rank order plots for each of the three age comparsions are shown. Comparisons are 30-month vs 10-month, 30-month vs 20-month, and 20-month vs 10-month. Each plot is ranked by normalized enrichment score.

Figure S2: Mean-difference (MD) plots of transposable elements (TE) expression Log2 FC against the average Log2 count-per-million (CPM) for the 30m/10m comparison (Fig 1D), 30m/20m comparison, and 20m/10m comparison. TE callouts list the: “TE name (**TE class**).”

Figure S3: Dendrogram and heatmap displaying the correlation between the 30 modules detected by WGCNA. The heatmap shows the Pearson correlation from -1 to 1 between each of the modules graphed along the x and y axes.

Figure S4: Expression of the canonical PGC-1a1 and alternative PGC-1a4 isoforms during neural stem cell (NSC) differentiation (n=3). Expression of the transcripts was determined by RT-qPCR. Significance was determined by one-way ANOVA.

Figure S5: Representative images (left)and distributions (right) of NAD(P)H fluorescence lifetime images of primary neurons and primary astrocytes. Representative images are artificially colored to show differences in mean fluorescence lifetime, the short component of the decay curve (τ_1_), the long component of the decay curve(τ_2_), or the relative contribution of free NAD(P)H (a1) to the mean fluorescence lifetime (n=10-12).

Figure S6: RT-qPCR detection of PGC-1a transcripts in control and LiCl-treated P1 astrocytes (n=6). Statistical significance was determined by Student’s t-test.

Figure S7: Immunohistochemical detection of GSK3β in the indicated hippocampal regions for 10-month-old, 20-month-old, and 30-month-old mice (n=4-6). Statistical significance was determined by 2-way ANOVA.
