## Supplementary figures and images for "Mitochondrial regulator PGC-1a in neuronal metabolism and brain aging"

### Figure S1

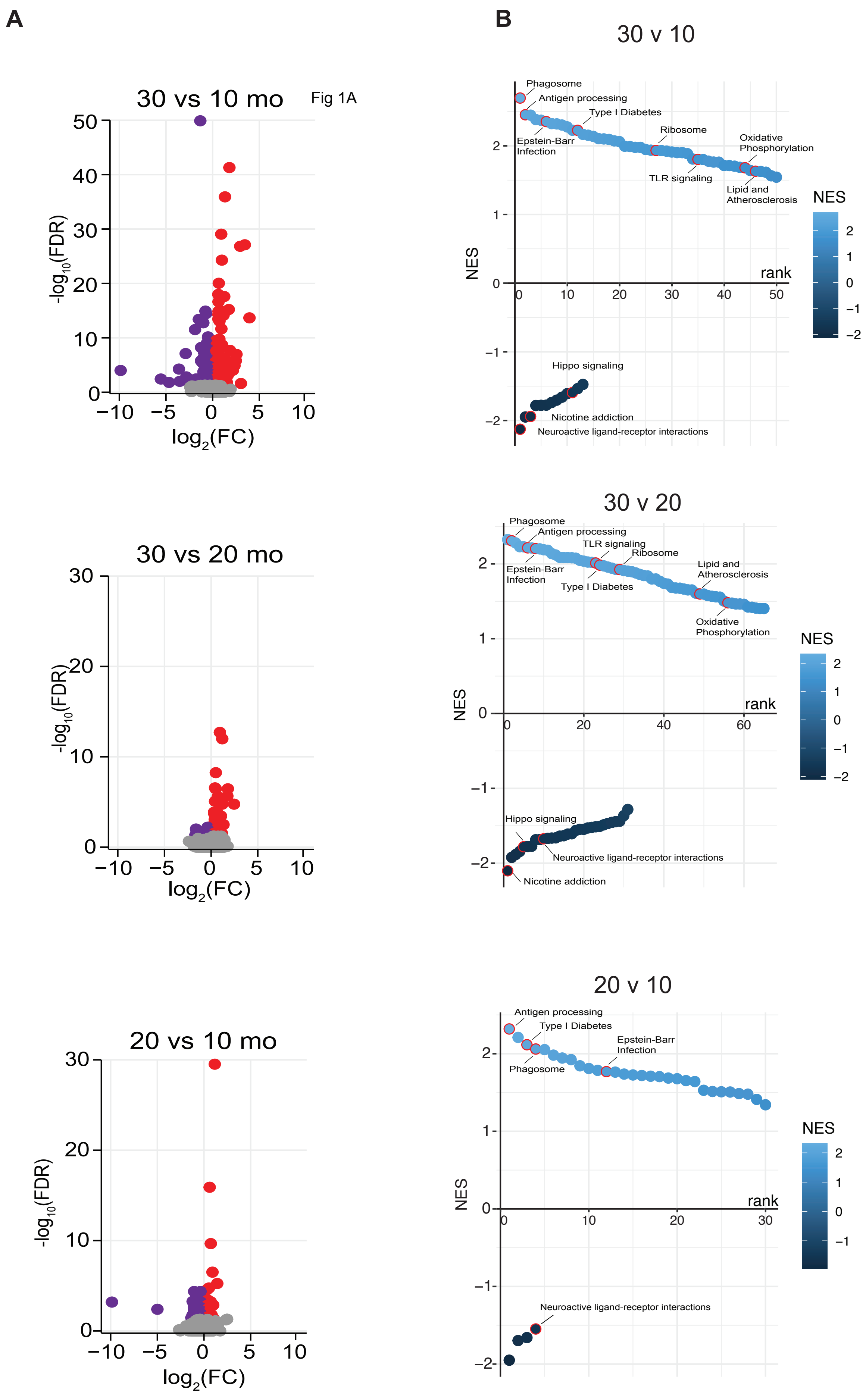

### Figure S2

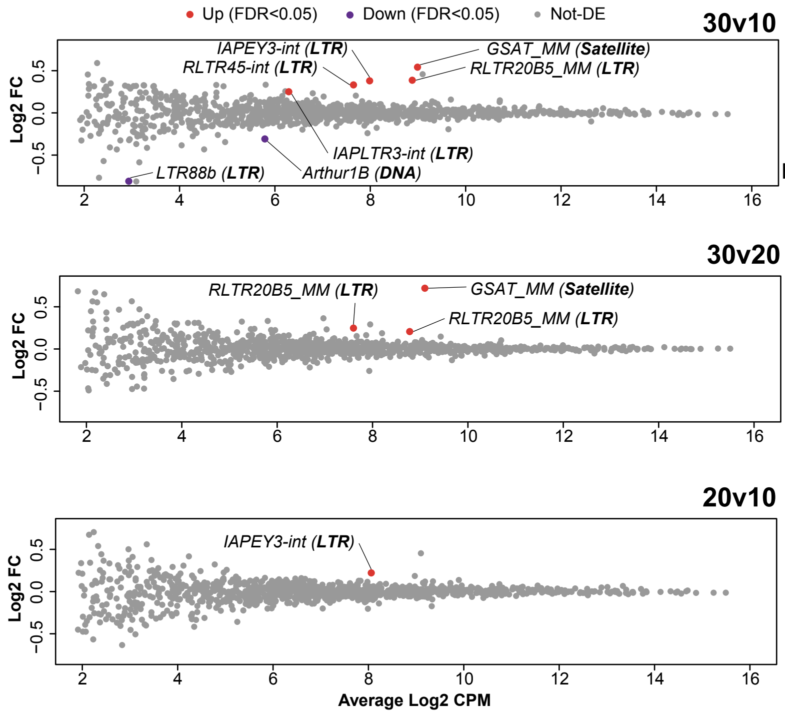

### Figure S3

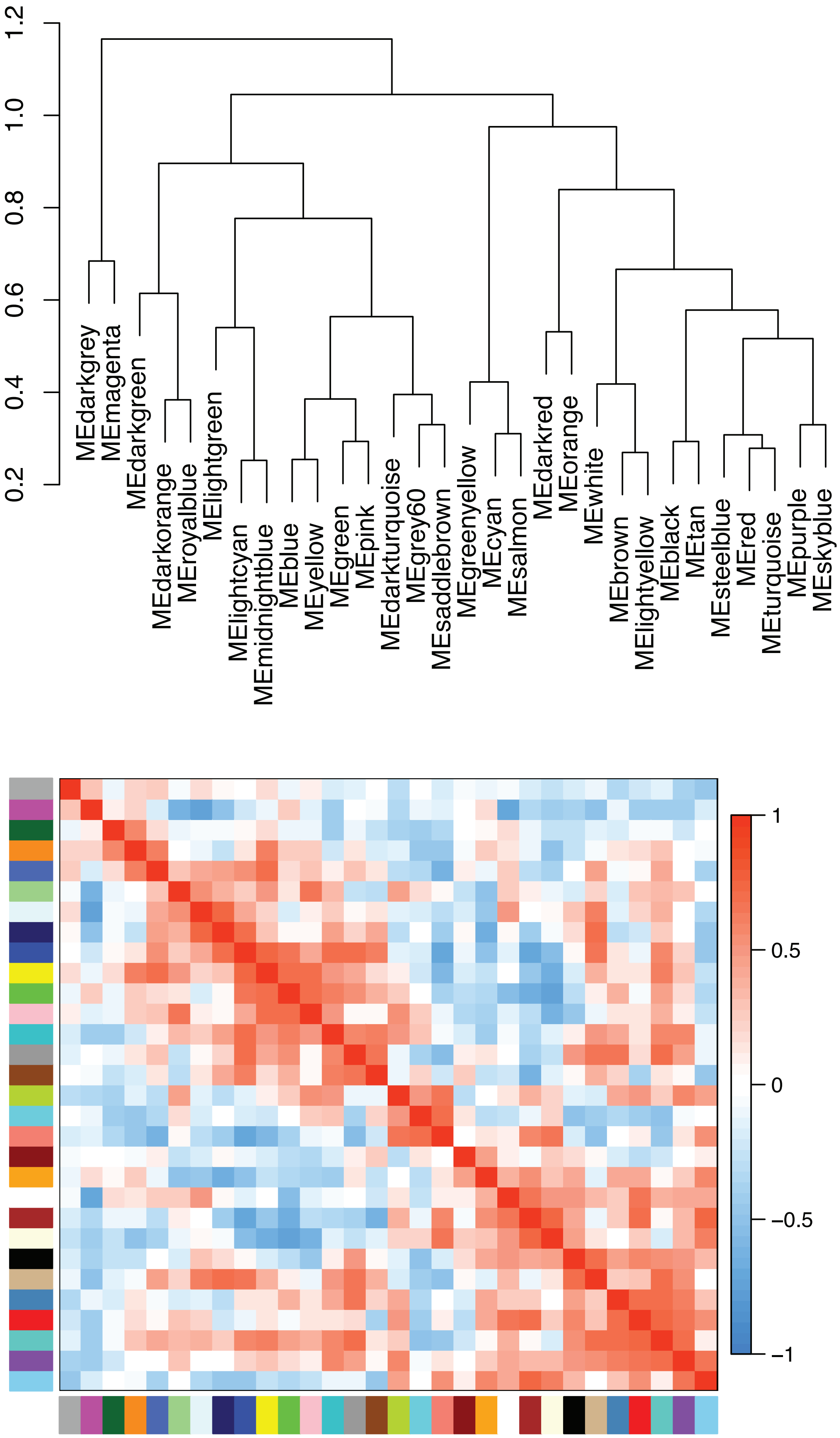

### Figure S4

##### NSC Differentiation

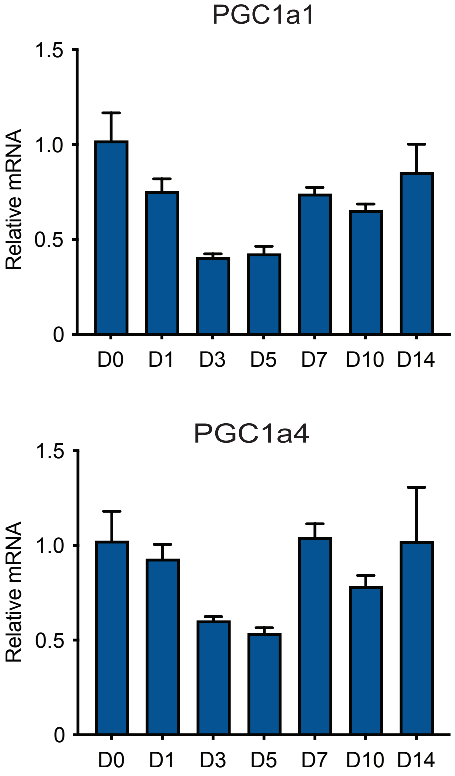

### Figure S5

#### Supplemental Figure: Neuron v. Astrocyte: MPLSM-FLIM Analysis

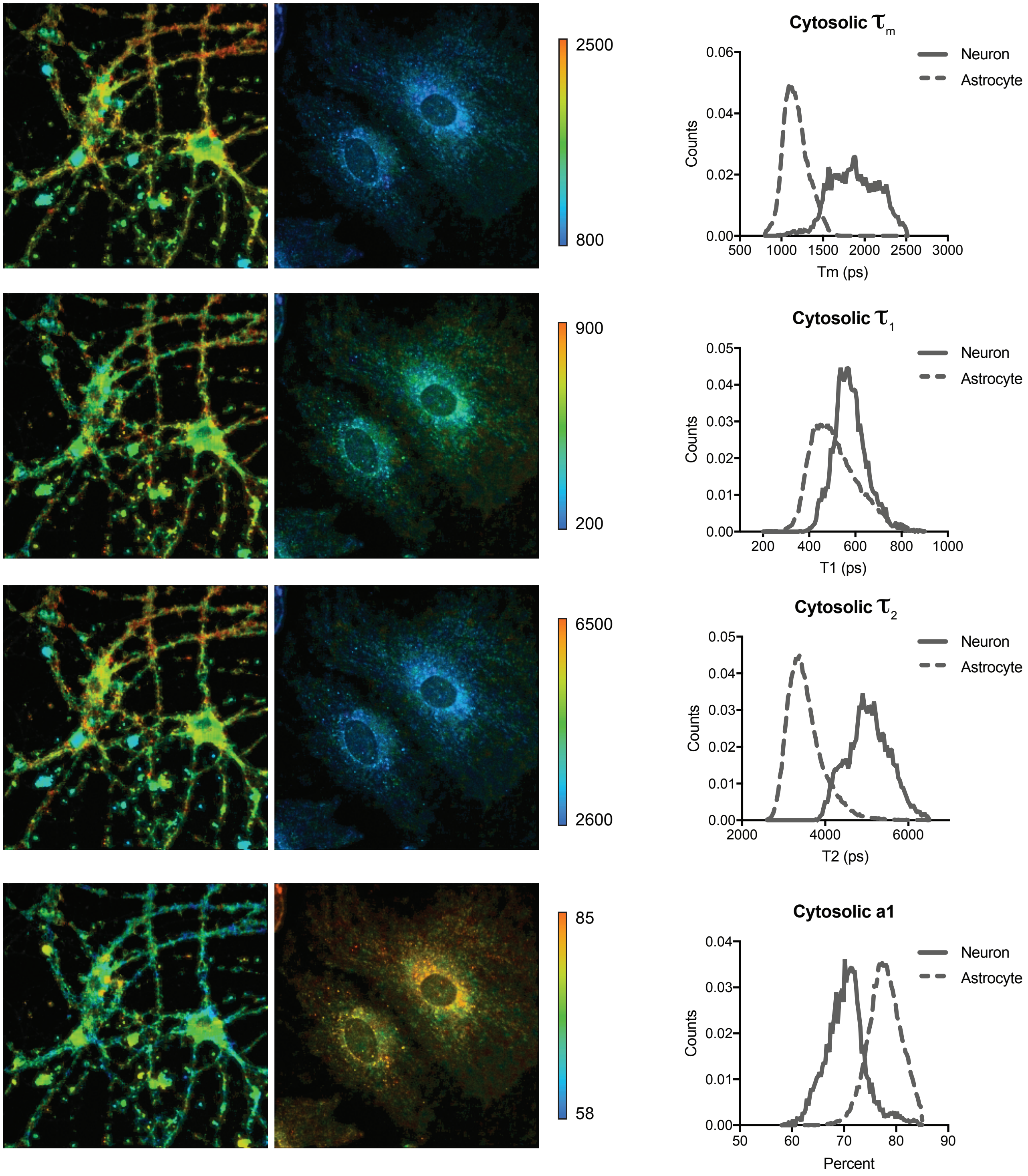

### Figure S6

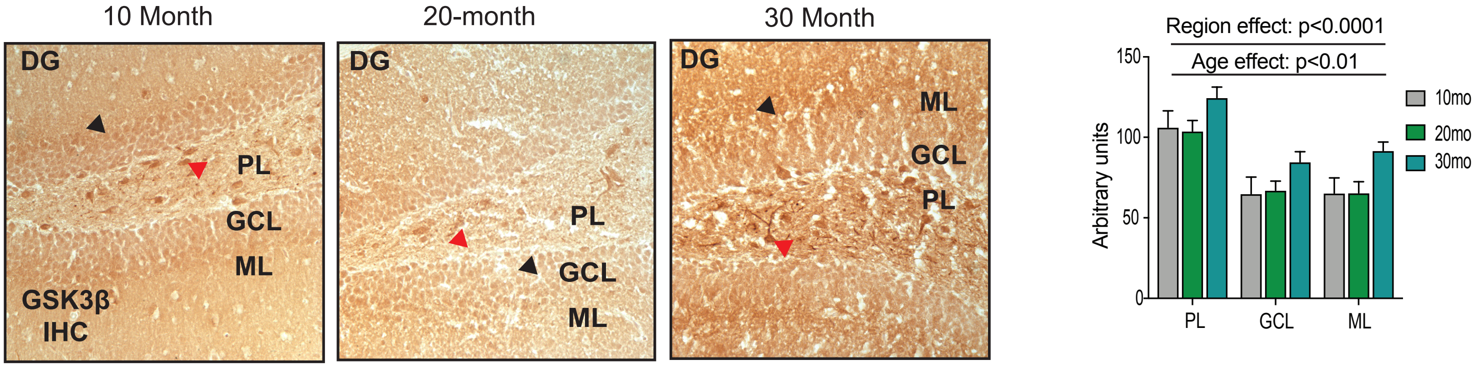

### Figure S7

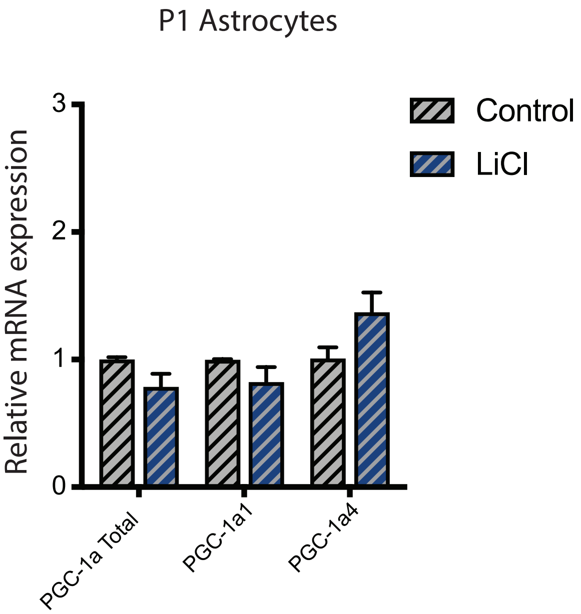
