## Supplemental Table Legends for "Mitochondrial regulator PGC-1a in neuronal metabolism and brain aging"

Supplementary Table 1: Differentially expressed genes and enriched KEGG pathways detected by GSEA of aged mouse brains.

Supplementary Table 2: Modules produced by WGCNA of aged mouse brains.

Supplementary Table 3: RT-qPCR primer sequences for PGC-1a transcript variants.

Supplementary Table 4: Differentially expressed proteins and significantly enriched KEGG pathways detected by overrepresentation analysis in aged mouse brains.

Supplementary Table 5: Differentially expressed genes and significantly enriched pathways detected by gene set enrichment analysis in LiCl-treated neurons.

Supplementary Table 6: Differentially expressed genes and significantly enriched pathways detected in brains of LiCO3-fed mice.
